# Loss of the mitochondrial carrier, *SLC25A1,* during embryogenesis induces a unique senescence program controlled by p53

**DOI:** 10.1101/2023.07.18.549409

**Authors:** Anna Kasprzyk-Pawelec, Mingjun Tan, Raneen Rahhal, Alec McIntosh, Harvey Fernandez, Rami Mosaoa, Lei Jiang, Gray W. Pearson, Eric Glasgow, Jerry Vockley, Christopher Albanese, Maria Laura Avantaggiati

**Author notes:** Correspondence: Maria Laura Avantaggiati.

## Abstract

Germline inactivating mutations of the SLC25A1 gene contribute to various human developmental disorders, including combined D/L-2-hydroxyglutaric aciduria (D/L-2HGA), a severe systemic syndrome characterized by the accumulation of both enantiomers of 2-hydroxyglutaric acid (2HG). The mechanisms by which SLC25A1 deficiency leads to this disease and the role of 2HG are unclear and no therapies exist. We now show that mice lacking both Slc25a1 alleles display a spectrum of alterations that resemble human D/L-2HGA. Mechanistically, SLC25A1 loss results in a proliferation defect and activates two distinct senescence pathways, oncogene-induced senescence (OIS) and mitochondrial dysfunction-induced senescence (MiDAS), both involving the p53 tumor suppressor and driven by two discernible signals: the accumulation of 2HG, inducing OIS, and mitochondrial dysfunction, triggering MiDAS. Inhibiting these senescence programs or blocking p53 activity reverses the growth defect caused by SLC25A1 dysfunction and restores proliferation. These findings reveal novel pathogenic roles of senescence in human disorders and suggest potential strategies to correct the molecular alterations caused by SLC25A1 loss.

## INTRODUCTION

The solute carrier family member SLC25A1, also known as CTP/CIC, belongs to a family of nuclear encoded ion transporters localized in the inner mitochondrial membrane where it promotes the bi-directional exchange of citrate between the mitochondria and the cytosol (Mosaoa et al, 2021; Palmieri et al, 2020; Ruprecht and Kunji, 2020). In the cytoplasm, citrate is a substrate for *de novo* lipid synthesis (DNL), while in the mitochondria it feeds into the tricarboxylic acid (TCA) cycle. The human *SLC25A1* gene maps to chromosome *22q11.2*. Heterozygous microdeletions of this region give raise to Velocardiofacial (VCFS) and DiGeorge (DGS) syndromes (Funato, 2022; Maynard et al, 2008), a group of developmental disorders associated with loss of 30 to 40 genes in addition to *SLC25A1*. Patients with *22q11.2* deletion syndrome present with heart malformations, cleft palate, immune deficiency due to thymic aplasia, short stature and abnormal facial features, particularly microcephaly, malposition of the ears, small eyes, underdeveloped chin and flat midface. We and others have previously shown that the knock-down of the *slc25a1a* homologous gene in zebrafish embryos induces craniofacial and heart abnormalities, suggesting a role for SLC25A1 in the pathogenesis of *22q11.2* deletion syndromes (Catalina-Rodriguez et al, 2012; Chaouch et al, 2014).

Various *SLC25A1* gene mutations have also been reported in a group of developmental disorders characterized by D/L-hydroxyglutaric aciduria (D/L-2HGA), a disease hallmarked by the accumulation and urinary escretion of the D- and L-enantiomeric forms of 2-hydroxyglutarate (2HG) (Pop et al, 2018; Edvardson et al, 2016; Prasun et al, 2015; Nota et al, 2015; Smith et al, 2013; Kranendijk et al, 2012). The clinical spectrum of manifestations associated with *SLC25A1* gene mutations in D/L-2HGA is very severe, consisting of a variety of craniofacial abnormalities, including facial dysmorphic features, down-slanting of the eyes, ear abnormalities, micrognathia or retrognathia, flat nose and microcephaly or macrocephaly (Pop et al, 2018). In addition, these patients present with developmental delay, brain abnormalities, (cerebral atrophy, agenesis of the corpus callosum), and with respiratory insufficiency, epilepsy, encephalopathy and occasionally, cardiomyopathy (Kranendijk et al, 2012). Unlike in VCFS/DGS, where only one allele of the *SLC25A1* gene is lost, D/L-2HG-aciduria results from homozygous or compound heterozygous mutations that typically involve a profound disruption of the citrate transport activity. We and others have previously reported that the most severe cases of D/L-2HGA are sustained by the compound heterozygous alleles, p.Ala9Profs*82 and p.Pro45Leu (onset at 1 day, death at 1 month) (Prasun et al, 2015;, Pop et al, 2018), which results in complete loss of SLC25A1 mitochondrial activities. In all other cases, with few exceptions, there is a strong correlation between the extent of loss of export activity and the severity of the clinical manifestations.

It is currently unknown whether 2HG plays a direct role in the pathogenesis of SLC25A1-associated D/L-2HGA. Both enantiomers are derived from the reduction of 2-oxoglutarate (α-ketoglutarate, αKG) and there are different pathways for 2HG production and elimination (Fig. 1A) (Du et al., 2022; Ježek 2020; Losman and Kaelin, 2013). Individual forms of L- or D-2HGA are caused by mutations in the 2-hydroxyglutarate dehydrogenases L-(*L2HGDH*), or D- (*D2HGDH*) which eliminate each enantiomer by oxidation to αKG. Gain of function, mutant forms of *isocitrate dehydrogenase 1 or 2* (*IDH1*/2) convert αKG to D-2HG, and *IDH2* mutations can also cause D-2HGA (Losman and Kaelin, 2013; Kranendijk et al, 2012). Mechanistically, D/L-2HG have been extensively characterized in the context of tumors harboring genetic mutations of *IDH1/2* (Ježek, 2020). However, these gain of function proteins produce extraordinarily high (10-30mM) amounts of 2HG. By contrast, it was initially noted that while combined D/L-2HGA is clinically more severe, the levels of D-2HG and L-2HG in these patients are very moderately elevated compared to other forms of 2HGA and this was considered a distinct feature of D/L-2HGA (Kranendijk et al, 2012). Furthermore, distinct biochemical alterations clearly separate D/L-2HGA sustained by SLC25A1 deficiency from other forms of 2HGA, particularly the presence of increased lactate and TCA cycle intermediates in the urine, indicative of mitochondrial dysfunction (Smith et al, 2016; Cohen et al, 2018; Prasun et al, 2015). Indeed, we have documented that genetic or pharmacologic inhibition of SLC25A1 in tumor cells impairs mitochondrial oxidative phosphorylation (OXPHOS) and rewires the metabolism towards glycolysis, essentially driving a Warburg effect (Mosaoa et al, 2021; Fernandez et al, 2018). However, whether and how SLC25A1 dysfunction influences mitochondrial activity in human diseases is currently uknown.

**Figure 1.**
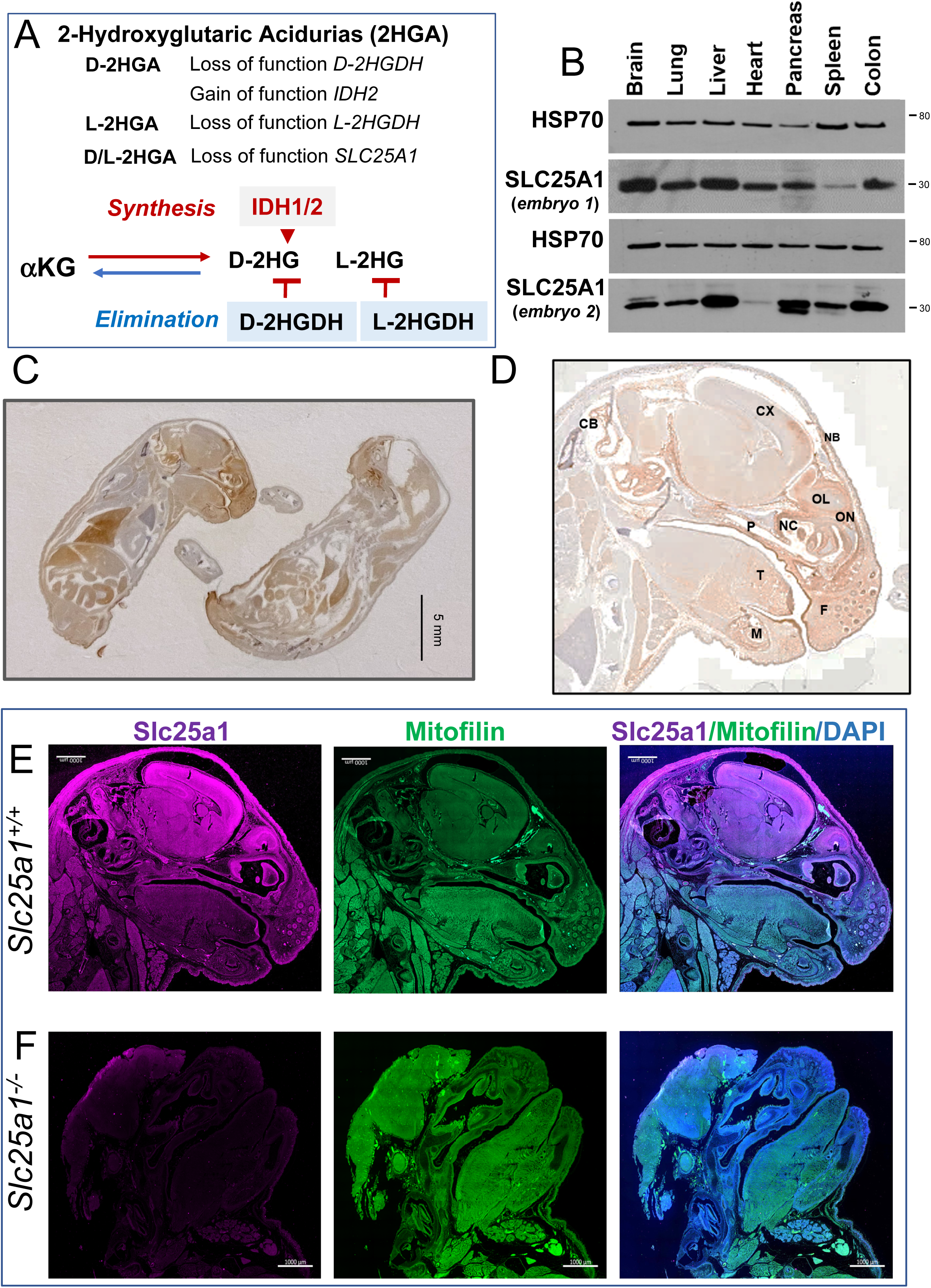
Analysis of SLC25A1 protein expression. (**A**) Schematic representation of 2-Hydroxyglutaric acidurias (2HGA). Upper panel: genetic alterations in individual and combined 2-HGAs. Lower panel: main pathways to the synthesis and elimination of D- and L-2HG. (**B**) Expression levels of SLC25A1 protein in the indicated organs derived from two *Slc25a1* wild-type embryos at E19.5 dpf. (**C-D**) Representative IHC images with the anti-SLC25A1 antibody staining in embryos at E19 dpf. Cb= Cerebellum, Cx= Cerebral cortex, F=follicle, P=palate, T=tongue, M=mandible, NB= nasal bone, NC= nasal cavity, OL=olfactory lobe, ON=olfactory nerve. (**E**) Staining with anti-SLC25A1, anti-mitofilin and DAPI in embryos at E19 dpf. Bar=1000 µm. **(F)** Staining as in E in E19 dpf *SLC25A1* nullizygous embryos.

In keeping with the activity of SLC25A1 in the transport of citrate, the current model envisions that a deficit in the cytosolic citrate pool hampers *de novo* lipid synthesis (DNL). Indeed, the only therapy that has been so far attempted in D/L-2HGA patients consists of oral administration of citrate, but the prognosis of D/L-2HGA is poor and the effects of citrate complementation remain controversial (Phua et al, 2024). In the present work we now show that mice lacking both *Slc25a1* alleles exhibit several phenotypic characteristics typical of VCFS and DGS as well as of D/L-2HGA. Mechanistically, loss of SLC25A1 activity leads to a severe deficit of proliferative capacity and uniquely triggers two distinct pathways of cellular senescence, mitochondrial dysfunction associated senescence (MiDAS) and oncogene induced senescence (OIS). We identify several potential corrective strategies to reverse these senescence programs, potentially opening novel strategies for therapeutic intervention in these diseases.

## RESULTS

### Loss of the *Slc25a1* gene during embryogenesis disrupts multiple organs development

To understand the role of SLC25A1, we began by studying the pattern of expression of SLC25A1 protein in normal murine embryonic tissues. The analysis of different organs of *Slc25a1* wild-type embryos at 19.5 days post-fertilization (dpf) by immunoblot showed strong expression in the brain, heart, lung, liver, colon spleen and pancreas (Fig. 1B). This finding was confirmed with immunohistochemistry (Fig. 1C,D). Immunofluorescence of the head showed specific enrichment of SLC25A1 protein in the mitochondria, as revealed by staining with mitofilin (Fig. 1E,F). Within the craniofacial region, SLC25A1 signal was particularly strong in the frontal region, in the mouth, eyes, palate and tongue, as well as in the brain and cerebellum, all sites affected in human diseases sustained by its deficiency (Fig. 1C-E).

To understand the impact of SLC25A1 activity, we generated a knock-out mouse model. Mice harboring a heterozygous germline targeted deletion in one *Slc25a1* allele were produced by the Mutant Mouse Resource & Research Centers (MMRRC, C57BL/6-tm1a) with a targeting vector which is based upon the ‘knockout-first’ allele, as we have described previously (Tan et al, 2020). In this vector, an *IRES:lacZ* trapping cassette and a *floxed* promoter-driven neo cassette is inserted between intron *1* and *2* of the *Slc25a1* gene on chromosome 16 (EV1A,B). We first expanded a population of heterozygous *Slc25a1*^+/-^ mice, and intercrosses of these animals subsequently allowed studying the population of homozygous mutants. These null mice completely lacked SLC25A1 protein and mRNA in all tissues examined (Fig. 2A and EV1C,D). Relative to wild-type and heterozygous animals, *Slc25a1*^-/-^ null mice were detected at roughly normal Mendelian ratio (Fig. 2B). However, only few of them were born, succumbing immediately after birth (Fig. 2C). Importantly, these embryos were be detected at all stages of development from day 7 to day 19-21 dpf (EV1), indicating that the knock-out of the *Slc25a1* gene leads to a highly penetrant perinatal lethal phenotype, but is not overtly embryonic lethal. By contrast, *Slc25a1* heterozygous showed no obvious defects and normal lifespan, which we followed up to 1 year. The analysis of the *Slc25a1^-/-^* embryos at different stages of development revealed a substantial spectrum of alterations affecting multiple organs. The majority of them displayed small body size and anemia/pallor (Fig. 2D-G), malformation/absence of the ears, absence of one or both eyes, as well as cleft palate (Fig. 2E-G). The cranio-facial region and the brain were the most affected and 43% of embryos showed overt defects in neural tube closure, manifested by exencephaly or anencephaly. A repository of computed tomography (CT) images available through the mouse phenotype consortium at UC Davis (Groza et al, 2023; Dickinson et al, 2016), confirmed that the craniofacial region of *Slc25a1^-/-^*embryos was severely dysmorphic often with unrecognizable anatomic features, including bulging or absent eyes, dilated turbinates, enlarged and protruding tongue (Fig. 2H,I and EV2A). Brain and liver hemorrhage (EV2B), heart alterations, gastroschisis (Fig. 2H^3^ and EV2B), and bone abnormalities were also present. These latter were significant and include lack of skull (Fig. 2E and 2G), pectus excavatum (Fig. 2H^4^), abnormal curvatures and thickness of the spine and increased cartilage content in the vertebral column reminiscent of enchondromatosis (Fig. 2H^3^ and EV2CD). Interestingly, although we have not quantified these latter abnormalities, pectus excavatum, enchondromatosis and hemorrhages were previously described in mice harboring knock-in mutations of gain of function mutant *IDH1/2* alleles (Hirata et al, 2015; Sasaki et al, 2012). H&E staining and scanning electron microscopy (SEM) performed on the brain and craniofacial region revealed prominent disorganization in the developing neocortex of *Slc25a1^-/-^* embryos at E18 dpf, which showed diffused areas of nuclear pleomorphism with alterations in nuclear size, disorganization of the stroma, and mis-orientation of neurons in the distinctive cortical layers (Fig. 2I,J). Alterations in the mitochondrial morphology were also clearly detected (not shown). The comparison of chondroblasts and collagen from the nose cartilage revealed morphological differences, with an increase in cytoplasmic vacuolization seen in *Slc25a1* deficient animals, and disruption in the directionality of collagen fibers in the matrix (Fig. 2K). In addition, the tongue of virtually all *Slc25a1^-/-^* mice appeared abnormal in size and protruding, with thicker myocytes, altered directionality of fiber packages and increased connective tissue (Fig. 2I,L).

**Figure 2.**
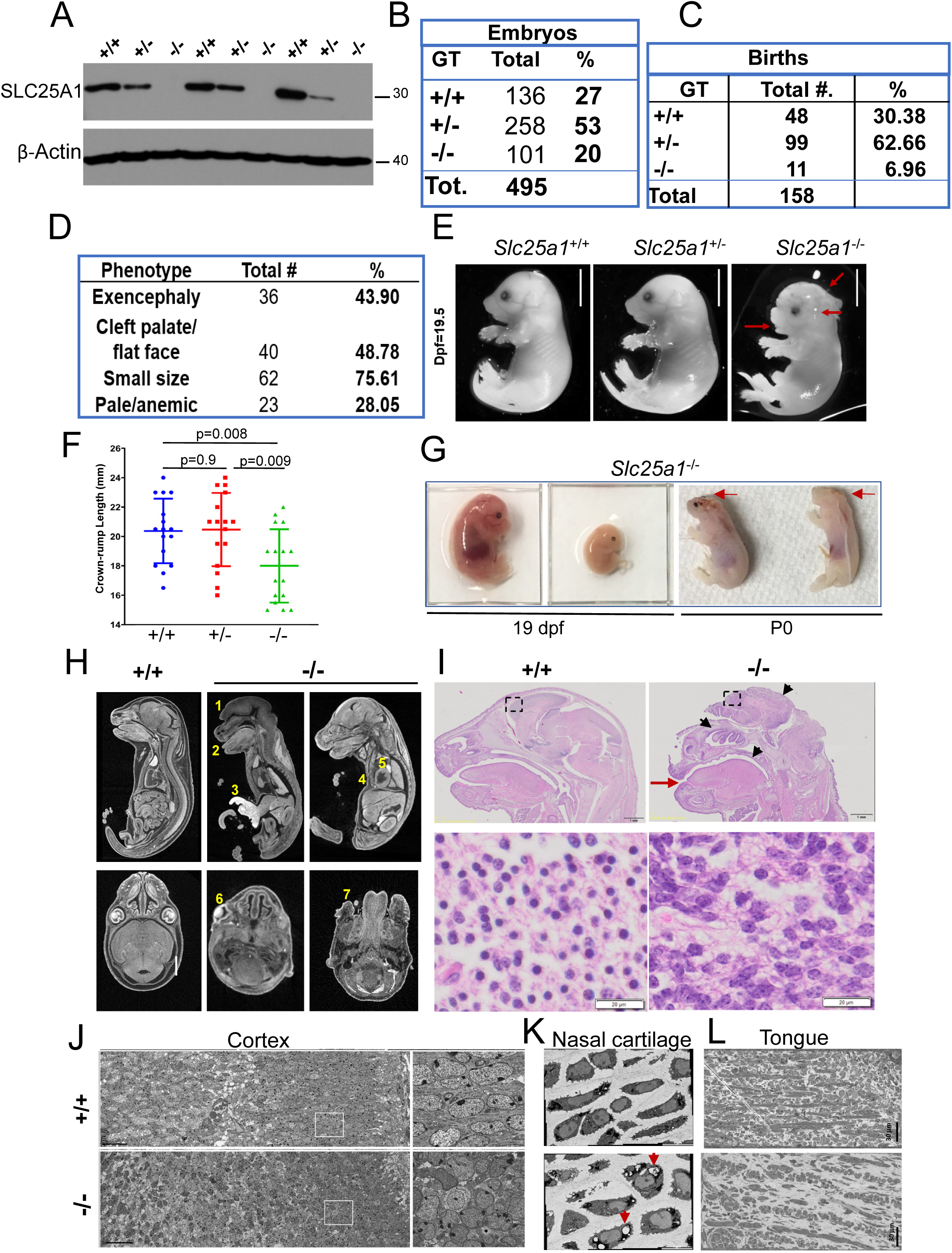
The knock-out of the *Slc25a1* gene leads to perinatal lethality and to multiple abnormalities. (**A)** Representative immunoblot showing the SLC25A1 protein expression in MEFs cells isolated from *Slc25a1*^+/+^, *Slc25a1*^+/-^ and *Slc25a1*^-/-^ embryos (n=3 per genotype). The antibody used in this blot is a polyclonal antibody raised against full length human SLC25A1, but other antibodies yielded similar results. (**B**) Representation of the genotypes observed across 85 litters (495 mice) obtained by breeding *Slc25a1* heterozygous mice. (**C**) Number and percentage of births observed across a population of newborn *Slc25a1*^-/-^ mice. (**D**) Prevalence of the phenotypic alterations observed in *Slc25a1*^-/-^ embryos. (**E**) Representative images of *Slc25a1*^+/+^, *Slc25a1*^+/-^ and *Slc25a1*^-/-^ embryos under stereoscopic microscope at E18.5 dpf. Red arrows point to cleft palate, underdeveloped ears and brain abnormalities. Bar=5 mm. (**F**) Measurements of crows to rump length showing smaller body size in *Slc25a1* homozygous mice compared with wild-type and heterozygous littermates (n= 16 mice per genotype, respectively). p-values were calculated with two tailed non parametric t-test. (**G**) Heterogeneity of the phenotypes of *Slc25a1*^-/-^ embryos at 19pdf or at P0, showing variations in body size, lack of skull and eyes in the last two embryos on left (indicated by arrows). (**H**) Computed Tomography (CT) scan of wild-type or *Slc25a1*^-/-^ embryos at E18.5 dpf. Yellow numbers indicate: 1: Excencephaly; 2: Protruding tongue; 3: Gastroschisis; 4: Pectus excavatum; 5: Abnormal heart. 6: Abnormal eye; 7: Abnormal facial area morphology. Bar=1mm. (**I**) Representative images of H&E staining in the embryos of the indicated genotype. Top panels show the anatomy of the whole head; lower panels show magnification of the cortical region and nuclear abnormalities in *Slc25a1*^-/-^ mice. Bar= 20 µm. Red arrow points to hypertrophic tongue. (**J-L**) Scanning electron microscopy (SEM) overviews of the neocortex (**J**), nasal cartilage (**K**) and tongue (**L**) in wild-type and *Slc25a1*^-/-^ embryos at E18.5 dpf. Arrows in K point to vacuolization in the nasal cartilage.

Thus, systemic loss of the *SLC25A1* gene during embryonic development recapitulates but also severely expands the spectrum of alterations seen in human DGS/VCFS and in D/L-2HGA.

### SLC25A1 deficient embryos undergo metabolic and transcriptional rewiring

The current model envisions that the pathogenesis of diseases due to impaired SLC25A1 activity is sustained by the deficit of citrate export from the mitochondria to the cytosol, which results in impairment of lipids synthesis. We tested this model by performing targeted and untargeted metabolomic and lipidomic analyses, paralleled by immuno-blot experiments interrogating the main pathways involved in lipid biosynthesis. The levels of the key rate-limiting, lipogenic enzymes, fatty acid synthase (FASN) and acetyl-CoA carboxylase Alpha (ACC1) were up-regulated in the brains and MEFs derived from *Slc25a1^-/-^* mice, at both the protein and mRNA levels (Fig. 3A-C). Given the herogeneity of the phenotypes of Slc25a1 deficient embryos and of the clinical manifestations of D/L2HGDH, for the metabolomic analyses we employed five wild-type and five *Slc25a1* null mice exhibiting different phenotypes, spanning from nearly normal to severe (EV2E). In spite of such heterogeneity, the principal component analysis (PCA) and the Partial Least Squares (PLS) revealed a good separation between these samples with the homozygous null mice clustering together (Fig.3D). Global metabolomic analysis showed that the main alteration consisted of enhanced triglyceride species in the brains and amniotic fluids (Fig. 3E,F), and enrichment in linoleic acid, phospholipids, glycerolipids, sphingolipids, arachidonic-, steroid-, and bile acid biosynthetic pathway, along with an increase in the Warburg effect (Fig. 3G). Importantly, the concentration of αKG, the precursor of 2HG, and of citrate were elevated *Slc25a1^-/-^* mice, and itaconate, which is derived from cytosolic citrate *via* the activity of Immune-Responsive Gene 1, *IRG1*, was increased (Fig. 3H), suggesting an excess of cytosolic citrate.

**Figure 3.**
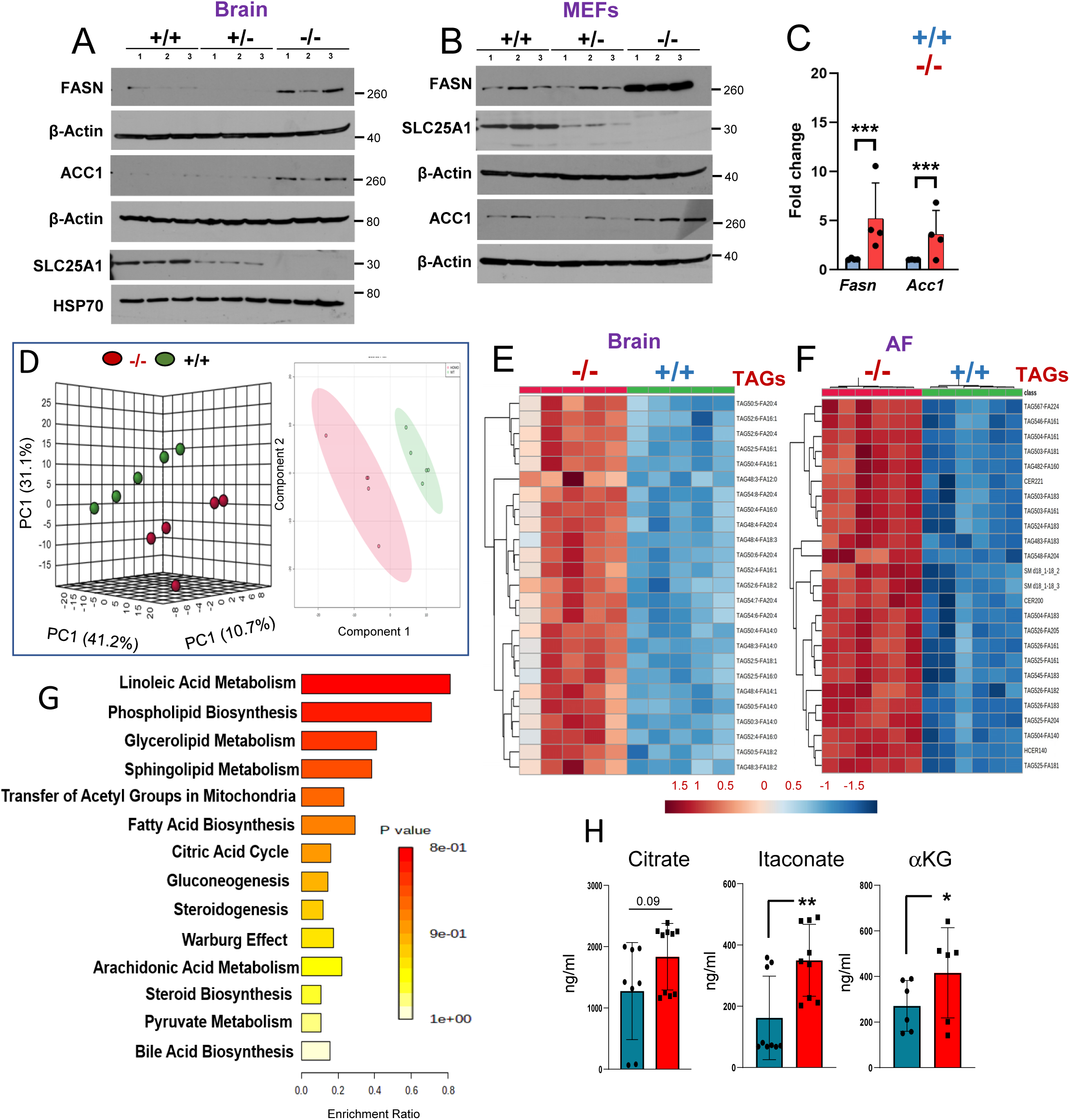
Loss of *Slc25a1 l*eads to metabolic remodeling. (**A-B**) Expression of the indicated proteins in brain (**A**) and MEFs (**B**) isolated from *Slc25a1*^+/+^, *Slc25a1*^+/-^ and *Slc25a1*^-/-^ embryos (n=3 per genotype). (**C**) mRNA levels detected with q-RT-PCR of the indicated genes in MEFs *Slc25a1*^+/+^ (blue) and *Slc25a1*^-/-^ (red). Each dot represents MEFs isolated from a different mouse. (**D**) Principal component analysis (PCA) scores plot and the partial least squares (PLS) of 5 wild-type and *Slc25a1* nullyzygous embryos used for metabolic studies. (**E-F**) Heatmap of top 25 enriched lipids in brain and AF samples obtained from *Slc25a1*^+/+^ (green) and *Slc25a1*^-/-^ (red) (n=5, from different litters, each in triplicate). Normalization was performed by dividing area under the curve for each metabolite by internal standard area. Clustering was performed by using the Metaboanalyst software. (**G**) Pathway enrichment analysis in amniotic fluid obtained from *Slc25a1*^-/-^ embryos relative to wild-type (n=3 per genotype, each in 3 technical replicates). (**H**) Concentrations of the indicated TCA intermediates in the amniotic fluid obtained from *Slc25a1*^+/+^ and *Slc25a1*^-/-^ embryos (n=3-5, each in technical replicates). All data are presented as mean value ± SD. Unpaired non parametric t-test was used throughout.

To then identify the pathways altered in an SLC25A1-dependent manner and particularly differences relative to the severity of alterations, we conducted a phenotype-to-transcriptome comparison using *Slc25a1^-/-^*mice with varying phenotypic alterations, classified as severe (E-SEV) or moderate (E-MOD) (Fig. 4A). E-SEV mice displayed critical abnormalities, including severe facial deformities like anophthalmia and cleft palate, transparent skin, along with brain alterations common in *Slc25a1* deficient embryos. Global transcriptome and hierarchical clustering identified 1395 genes uniquely regulated in *Slc25a1* homozygous mutants with most differentially regulated genes being up-regulated (Fig. 4B,C). These included: genes involved in inflammatory and pro-oncogenic pathways (Immune Cytokine Signaling, Signaling by NRTKs, AP1 Transcription & MAPK Signaling), the hypoxia response and, surprisingly, senescence (Fig. 4D), along with an enrichment of tumor-associated transcriptional signatures involved in breast cancer, neuroblastoma, ovarian cancer, and others, as well as induction of the p53 tumor suppressor (Fig. 4E). Using the transcriptomic data derived from each of the embryos (Fig. 4F), we then employed a Hue-Saturation-Value (HSV) analysis and the Enrichr gene set tool, to identify genes that are significantly more active in E-SEV and moderately elevated in E-MOD *versus* wild-type (*WT.**>** E-MD.**>>** E-SV);* p< 0.05) (Xie et al, 2021; Kuleshov et al, 2016). This approach revealed that the top up-regulated signals in the E-SEV are the hypoxia response, along with pro-inflammatory (TNFα, IL-6) and pro-oncogenic pathways, including mTORC, the KRas pathways, and the induction of the Epithelial to Mesenchymal Transition (EMT) (Fig. 4G). In addition, both glycolysis and cholesterol responses were enhanced, consistent with the previous metabolic analysis.

**Figure 4.**
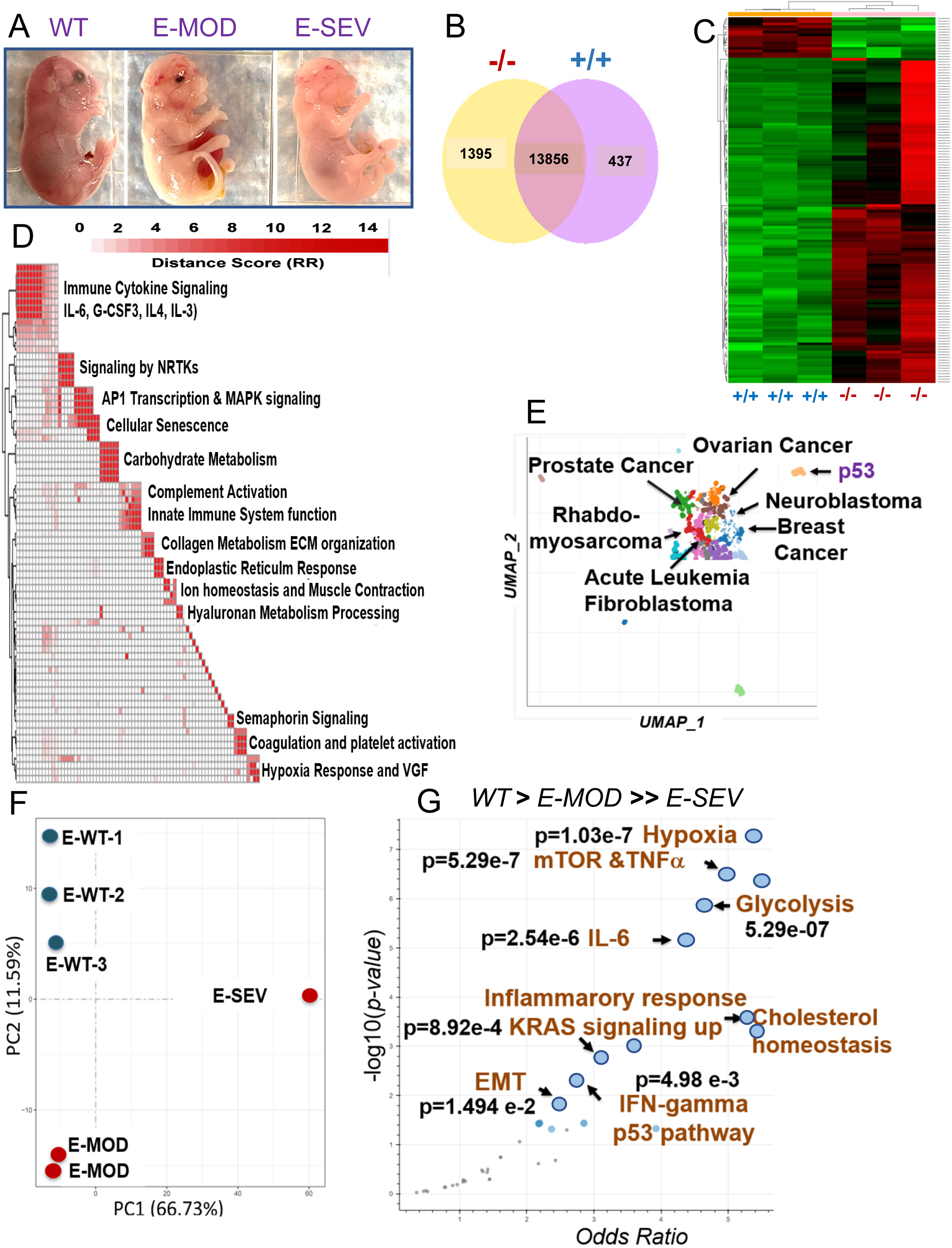
Transcriptional analysis of *Slc25a1* deficient embryos. (**A**) Representative phenotypes of *Slc25a1^-/-^*mice classified based on the severity of the phenotype (E18-19 dpf). (**B**) Venn diagram representing genes differentially regulated in the brains isolated from *Slc25a1*^+/+^ and *Slc25a1*^-/-^ embryos at 18.5 dpc (n=3). (**C**) Hierarchical clustering map of differentially expressed genes as in B. (**D**) Pathway enrichment analysis of genes significantly elevated in *Slc25a1*^-/-^ mice relative to wild-type. All significantly enriched pathways (FDR<0.05), assessed from the Reactome Database, were clustered *via* a distance metric derived from shared significant genes common amongst enriched gene sets. Clustered gene sets were summarized by biological function. (**E**) Scatter plot of enriched transcriptomic signature (with ChEA). (p=<0.05). (**F**) PCA plot of the transcriptomic profiles of indicated embryos. (**G**) Transcriptomic data derived from each of the embryos were used for a Hue-Saturation-Value (HSV) analysis. The Enrichr gene set tool was employed to identify gene sets that are significantly more active in E-SEV and moderately elevated in E-MOD *versus* wild-type (*WT.**>** E-MD **>>** E-SV);* p< 0.05.

We conclude that in this mouse model, SLC25A1 is not rate limiting for maintenance of the citrate and lipid pool during embryonic development as generally thought, and consistent with the well documented existence of multiple pathways able to replenish cytosolic citrate in cells lacking SLC25A1, including *via* glutamine reductive carboxylation (Mosaoa et al, 2021; Jiang et al, 2017; Jiang et al, 2016;). Instead, SLC25A1 loss rewires the transcriptome towards senescence, pro-oncogenic and inflammatory signals and induces an hypoxic response.

### SLC25A1 loss induces the IOS and MiDAS pathways of senescence

We next examined the growth properties of embryo fibroblasts (MEFs) derived from mice, as well as of clinically relevant samples, using skin fibroblasts from two newborn patients affected by D/L-2HG-aciduria, patient 893 (male) and patient 897 (female) (Phua et al, 2024). Both the *Slc25a1^-/-^*MEFs cells and the patient-derived fibroblasts exhibited a proliferation defect at early passages (Fig. 5A and not shown). This loss of proliferative capacity was highly suggestive of premature senescence, which when aberrantly induced during embryonic development drives neuro-developmental and neural tube closure defects (Klein et al, 2023; Rhinn et al, 2022; Xu et al, 2021). Consistent with the transcriptomic analysis and with the idea that loss of Slc25a1 induces senescence, the *Slc25a1^-/-^*MEFs as well the patients fibroblasts showed a strong senescence-associated beta-galactosidase (SAβ*-*GAL) signal, the typical hallmark of senescent cells (Fig. 5B-E). Further, the head and brain of *Slc25a1^-/-^* mice harvested at late development stage (E19 dpf), reacted with antibodies directed against p21, a key mediator of senescence (Fig. 5F,G). To rule out that senescence is the result of passaging primary cells in culture and and to establish the relevance of the Slc25a1 transport activity, we conducted acute SLC25A1 knock-down experiments, or we employed the specific SLC25A1 ihibitor (SLC25A1-i) (Mosaoa et al, 2021). The shRNA, as well as the SLC25A1-i induced SAβ*-*GAL activity and arrest at the G1/G2 phase of the cell cycle (Fig. 5H,I), demonstrating a direct effect of SLC25A1 inhibition in the induction of the senescence arrest.

**Figure 5.**
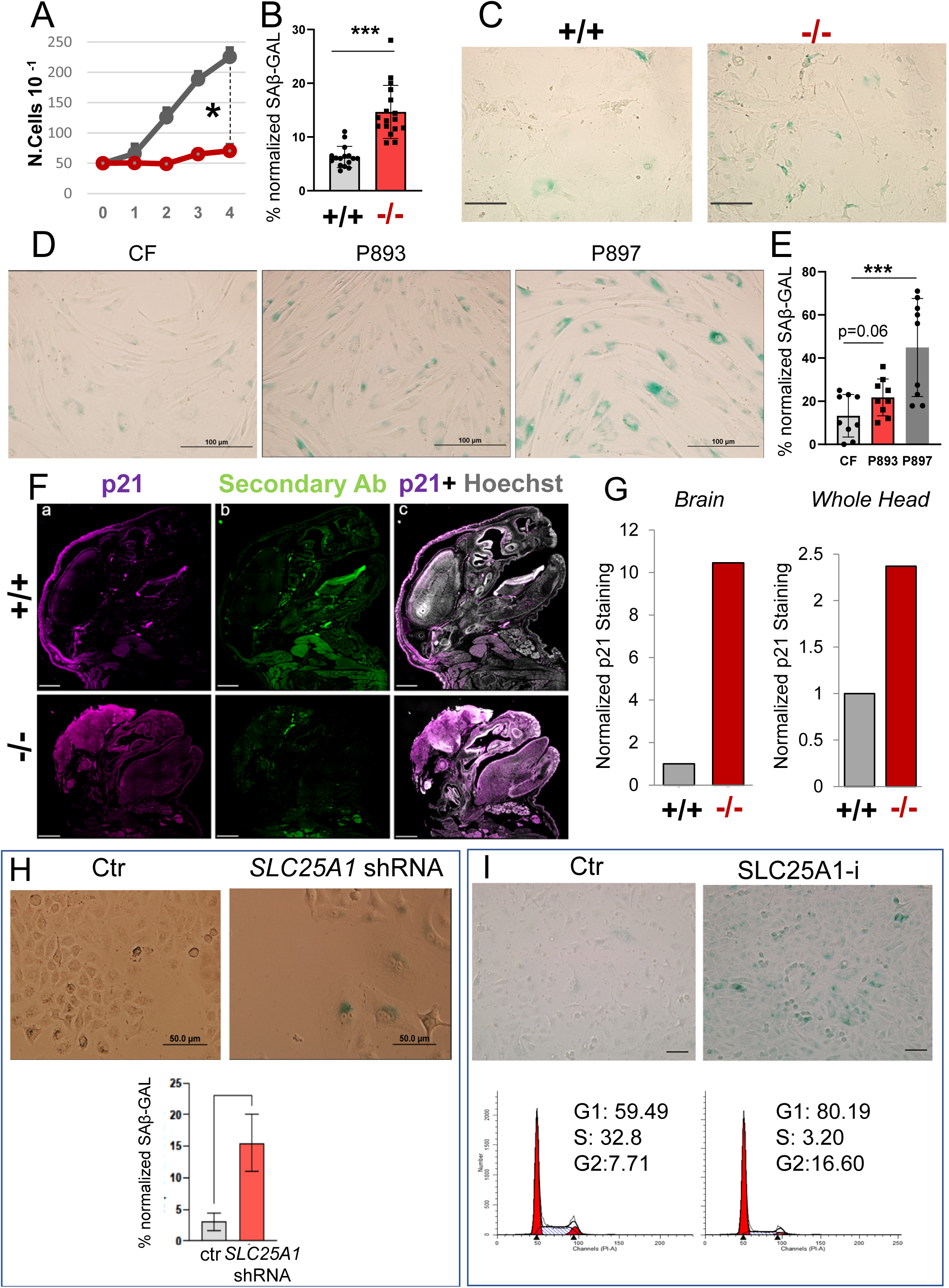
Induction of premature senescence in cells and tissues lacking SLC25A1. (**A**) Growth curves of MEF cells isolated from *Slc25a1*^+/+^ and *Slc25a1*^-/-^ embryos. (**B-C**) Quantification of the SA-β*-*GAL positive MEF cells (%) normalized to the total number of cells identified with DAPI counter-staining pooled from two independent experiments and representative images of (SA-β*-*GAL) activity. (**D**-**E)** Representative images of SA-β*-*GAL staining in human fibroblasts (**D**) and quantification of the SA-β*-*GAL positive cells normalized to the total number of cells identified with DAPI counter-staining. (**E**). All IF data are presented as a % positive cells ± SD pooled from two independent experiments. Bar=100 μm. (**F**) p21 immunofluorescence staining of the head of *Slc25a1*^+/+^ and *Slc25a1*^-/-^ embryos (E19 dpf). Hoechst was used to counterstain the nuclei. Secondary antibody alone was used as a control. Bar= 1000 µM. (**G**) Quantification of experiments shown in F with imageJ representing particle counts normalized to DAPI of the brain or whole head. (**H**) A549 cells were transduced with the control lentivirus or with lentivirus harboring the SLC25A1 specific shRNA (see Fig.10 for SLC25A1 levels), selected with puromycin for 5 days and stained with β*-*GAL. (**I**) SA-β*-*GAL activity and cell cycle analysis of A549 cells treated with the SLC25A1-i.

Senescent cells secrete many factors known as the Senescence-Associated Secretory Phenotype (SASP), which differ depending upon the upstream triggering signal(s) (Fig. 6A). Oncogene induced senescence (OIS), is characterized by secretion of pro-inflammatory mediators IL-6, IL-1β, and IL-1α, and involves NFkB, mTOR, AP1/MAPK signaling, and the JAK/STAT pathway (Schmitt et al., 2022; Wang et al., 2022; Cuollo et al., 2020; Xu et al., 2015). Due to DNA replication stress imposed by oncogenes, OIS is associated with the DNA damage response, hallmarked by induction of phosphorylated histone gamma H2AX and of the kinase Chk1/2 (Poehlmann A et al, 2011). A second form of senescence, mitochondrial dysfunction-induced senescence (MiDAS) contributes to accelerated aging in murine models of progeroid syndromes (Wile et al., 2016), and instead lacks pro-inflammatory cytokines while recruiting the activity of AMPk, TNFα, and of anti-inflammatory IL-10, among others (Wiley and Campisi, 2021; Wiley et al., 2016; Gomes et al., 2013). MiDAS occurs when the NAD+/NADH ratio drops, leading to mitochondrial dysfunction and activation of the hypoxia inducible factor, HIF1α in normoxic conditions, defined as a pseudo-hypoxia response. Both OIS and MiDAS activate p53 to enforce growth arrest.

**Figure 6.**
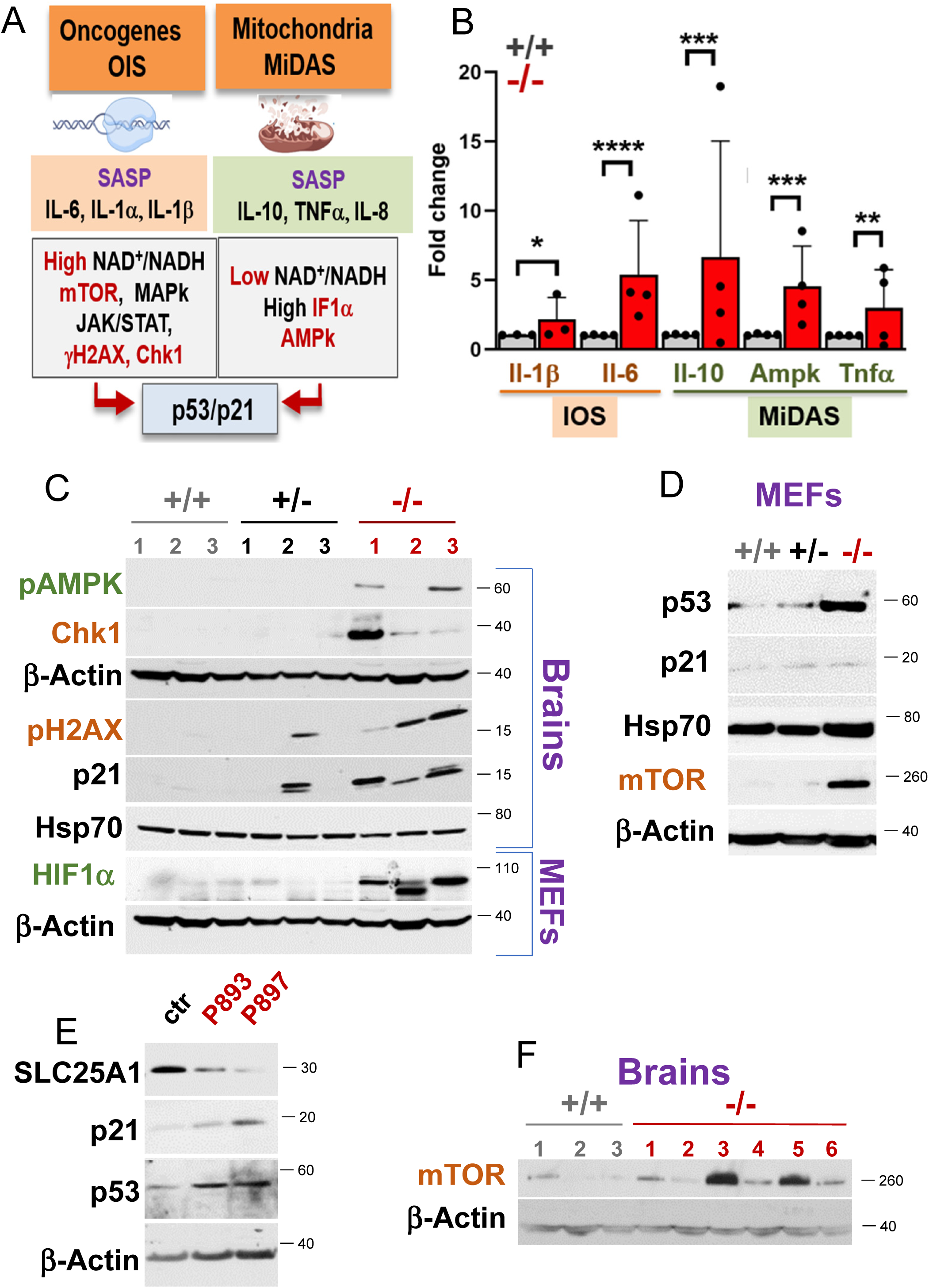
SLC25A1 dysfunction induces the OIS and MIDAS pathways of senescence. (**A**) Scheme of the OIS and MiDAS pathways of senescence and of the associated SASPs, depicting DNA replication stress or mitochondrial dysfunction, respectively (see also text for explanation) (*created with Biorender*). (**B**) mRNA levels of the IOS (brown) and MiDAS (green) SASPs in MEFs *Slc25a1*^+/+^ (grey) and *Slc25a1^-^*^/-^ (red) respectively (n=3-4). (**C-F**) Expression level of the indicated proteins involved in IOS (brown) and MiDAS (green) in the brain samples (**C,F**), MEFs (**C**,**D**) and in the patient’s fibroblasts (**E**) (n=1-6).

The analysis of the senescence program driven by SLC25A1 loss revealed a robust increase in IL-1β, IL-6, γH2AX, Chk1 and mTOR indicative of OIS, along with IL-10, TNFα and AMPk which are typical of MiDAS (Fig. 6B-D). Further, the levels of HIF1α were increased in the *Slc25a1^-/-^* MEFs cultured *in vitro* in normoxic conditions, suggestive of a pseudo-hypoxia response (Fig. 6C), and p53 and p21 were induced in both the *Slc25a1* deficient mice and in the patient-derived fibroblasts indicative of senescence arrest (Fig. 6C-E). Consistent with the previous transcriptomic data, the levels of mTORC, which plays a role in nutrient sensing and also as a mediator of senescence associated to aging (Weichhart, 2018), were elevated in most of these brains (Fig. 6F).

Collectively, these findings support the notion that loss of SLC25A1 leads to the simultaneous activation of both the IOS and MiDAS pathway. Given these results, we next investigated the molecular mechanisms by which SLC25A1 deficiency drives the MiDAS and IOS response.

### SLC25A1 dysfunction impairs OXPHOS *via* depletion of NAD^+^ and enhanced PDK1-mediated glycolysis

The NAD^+^/NADH ratio is a key determinant of OXPHOS. Several steps in the TCA cycle generate NADH from NAD+ and this chain of reactions is necessary for respiration *via* the activity of Complex-I (CI), which donates the electrons derived from NADH to other respiratory complex subunits within the electron transport chain (ETC). First, we determined that the *Slc25a1^-/-^* MEFs exhibited a severe deficit in the basal (BR), maximal (MR) and spare respiratory capacity (SRC) and of ATP production (Fig. 7A), coinciding with a significant loss of function of complex I, II and III and V (Fig. 7B). A similar respiratory deficit was recently described in the patient fibroblasts (Puah et al, 2024). In addition, cells lacking SLC25A1 prominently turned on glycolysis (Fig. 7C), which consumes NAD^+^ because many glycolytic enzymes use NAD^+^ as a co-factor (Xie et al, 2020). To then establish whether SLC25A1 dysfunction depletes NAD^+^ at least in part by turning on glycolysis, we again turned to a controlled cellular system where SLC25A1 activity was inhibited with the SLC25A1-i. By using an enzymatic activity assay that assesses the ability of lactate dehydrogenase-A, LDH-A, to carry out the oxidation of endogenous lactate to pyruvate, which reduces NAD^+^ to NADH, we found a stark 20 fold increase in the rates of conversion of NAD^+^ to NADH in cells treated with SLC25A1-i (Fig. 7D). Accordingly, the NAD^+^/NADH ratio was strongly reduced in the *Slc25a1^-/-^* MEFs (Fig. 7E).

**Figure 7.**
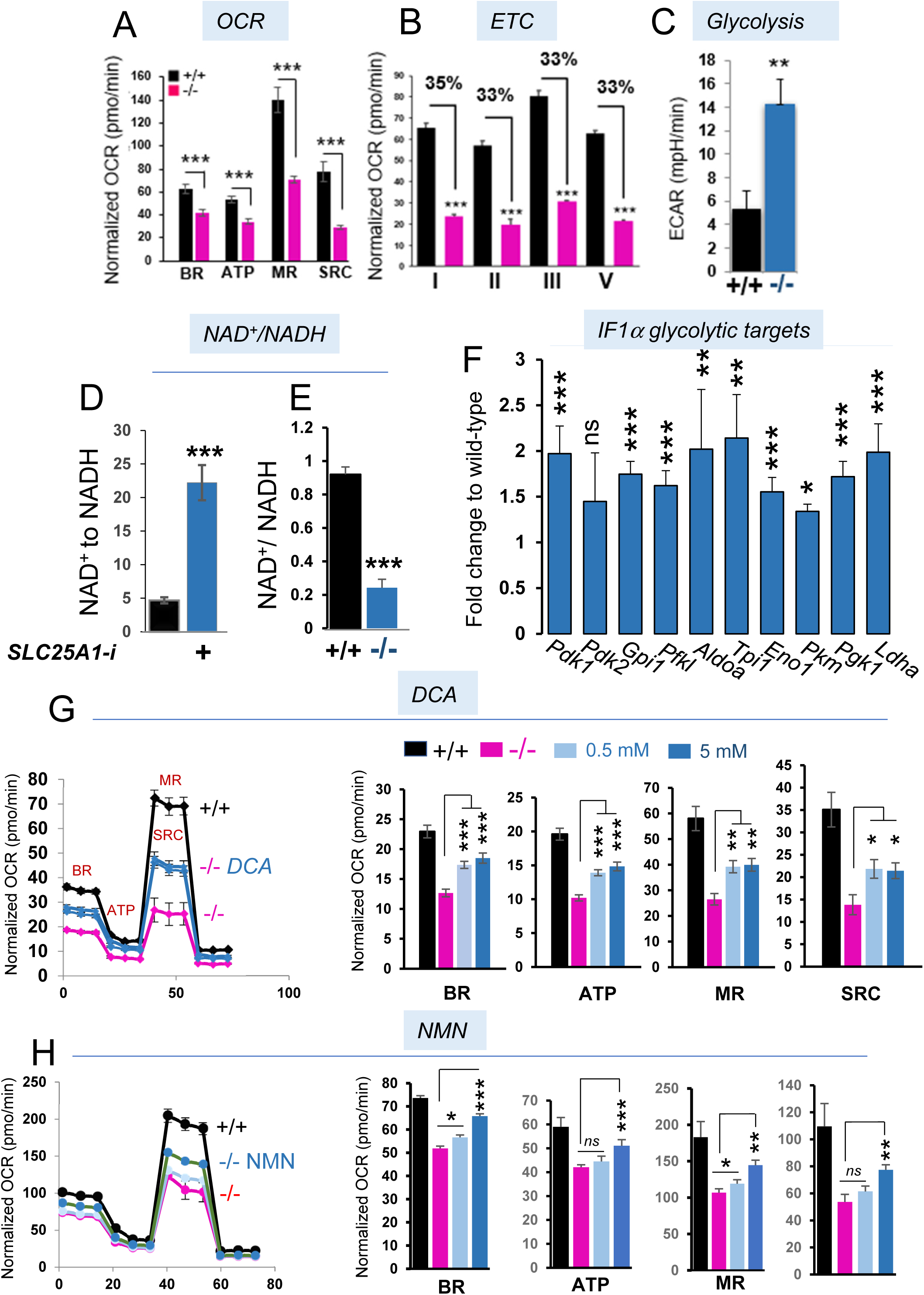
Cells lacking SLC25A1 undergo mitochondrial respiration deficit. (**A**) Oxygen consumption rate (OCR) was assessed using Seahorse Extracellular Flux Analyzer in MEFs isolated from *Slc25a1*^+/+^ (black) and *Slc25a1^-^*^/-^ (pink) embryos. Injection of Oligomycin, FCCP and Rotenone/Antimycin allowed calculation of the BR= Basal respiration; ATP= ATP Production, MR= Maximal Respiration, SRC= Spare Respiratory Capacity. (**B**) Activities of the indicated ETC complexes measured in the ^+/+^ or ^-/-^ MEFs. % = residual activity. (**C**) Glycolysis in the indicated group of MEFs measured as Extracellular acidification (ECAR). (**D**) Enzymatic activity assay using LDH-A in cell extracts derived from cells untreated or treated with the SLC25A1-i, and measured by absorbance at 340nm, as rate of conversion of NAD^+^ to NADH. (**E**) NAD^+^/NADH ratio in freshly isolated MEFs from *Slc25a1*^+/+^ and *Slc25a1^-^*^/-^ embryos. (**F**) Relative expression of indicated HIF1α genes/transcripts in brains isolated from *Slc25a1*^+/+^ and *Slc25a1^-^*^/-^ embryos. (**G-H**) OCR measured in *Slc25a1*^+/+^ and ^-/-^ MEFs untreated or treated with the indicated concentration of DCA (**G**) for 4 hrs, or NMN (**H**) for 16 hrs at 50 and 100 uM. Bars represent SD and p-values were calculated using two tailed non parametric t-test.

To identify the mechanisms leading to defective mitochondrial OXPHOS we focused on the pseudo-hypoxia response and on the glycolytic switch. Consistent with prior data, the brains of *Slc25a1^-/-^*mice were significantly enriched in glycolytic- and HIF1-signal pathways (Fig.4 and not shown), as well as in glycolytic genes that are under direct transcriptional control of HIF1α. These include the kinases PDK1 and PDK2, glucose transporters (MCT4, GLUT4), LDH-A, Enolase (ENO1), and glucose-phosphate-isomerase 1 (GPI1) (Fig. 7F). PDK1 inhibits the pyruvate dehydrogenase complex, PDH, preventing the conversion and entry of acetyl-CoA into the TCA cycle for OXPHOS, while enhancing its reduction to lactate. To then directly test the relevance of this pathway, we explored the activity of the PDK1 and glycolytic inhibitor dichloroacetate (DCA). As shown in Figure 7G, a short time treatment (4 hours) of the *Slc25a1^-/-^*MEFs with DCA significantly rescued the rates of OCR as well as of ATP production, to levels nearly comparable to wild-type MEFs. Similar results were obtained by supplementing cells with the cell permeable, NAD^+^ precursor Nicotinamide mononucleotide, NMN, which also partially rescued the rates of OXPHOS (Fig. 7H).

Albeit we cannot exclude that additional mechanisms are involved in the reduced OXPHOS rates driven by SLC25A1 deficiency, these data importantly demonstrate that the glycolytic switch is not simply a compensatory response to impaired mitochondrial respiratory activity, but is actually a driver of mitochondrial dysfunction and imply that such respiratory deficit can at least partially be corrected.

### In cells lacking SLC25A1 low levels of 2HG are produced by wild-type IDH1 and induce IOS

It was noted before that in combined D/L-2HG-aciduria caused by SLC25A1 impairment the concentrations of D-2HG and L-2HG are moderately increased compared to individual forms of L- or D-2HG-aciduria, which result from mutations of the *IDH1/2* or *D/L-2HGDH* genes and leads to high millimolar levels of these enantiomers (Ježek et al, 2020; Kranendijk et al, 2012). In agreement with these previous observations, we determined that the total pool of 2HG was only moderately elevated in the amniotic fluids of Slc25a1 deficient mice albeit being higher in the brains, reaching only a five-fold average enrichment compared to control (Fig. 8A,B). To then determine the relative enrichment of the D- and L-enantiomer, a chiral derivatization method was employed (Cheng et al, 2015) (see EV3A for the L/D peaks separation). We established that the D-enantiomer accounts for approximately 70% of the total pool and represents the main pool enriched in *Slc25a1^-/-^* mice relative to control (Fig. 8C,D). To further validate these results, we analyzed previously published data derived from nine patients affected by D/L-2HGA (Pop et al, 2018; Prasun et al, 2015), and found a comparable, albeit not esclusive prevalence of D-2HG (Fig. 8E). Importantly, in cells harboring wild-type *IDH1/2* genes, the acute SLC25A1 knock-down (KD) led to a four to five fold accumulation of D-2HG, again demonstrating that loss of SLC25A1 activity directly, but modestly induces enrichment of this metabolite (Fig. 8F). To corroborate this important point, we also comparatively studied the levels of D-2HG in fibrosarcoma cells, HT-1080, which harbor one of the most common *IDH1* mutations, IDH1^R132C^. This analysis confirmed the significantly lower enrichment of this enantiomer in cells harboring the SLC25A1-KD relatively to cells expressing gain of function IDH1 mutants (Fig. 8G).

**Figure 8.**
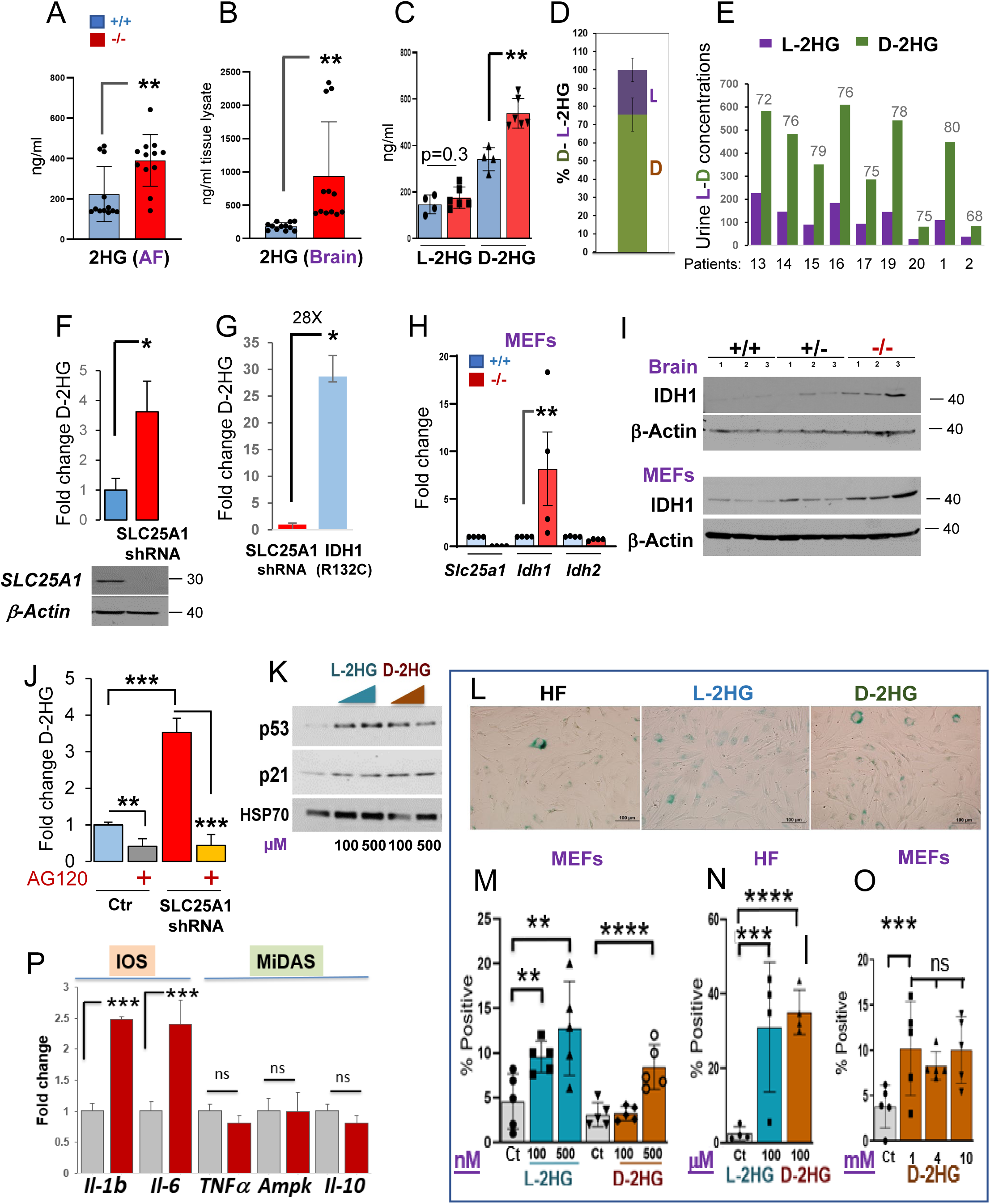
2HG induces the IOS pathway of senescence. (**A-B**) 2HG enrichment in the amniotic fluid (**A**) and brain samples (**B**) obtained from *Slc25a1*^+/+^ (blue) and *Slc25a1*^-/-^ (red) embryos at E18.5 dpf (n=4). (**C**) D-L- 2HG enrichment in the amniotic fluids of *Slc25a1*^+/+^ (blue) and *Slc25a1*^-/-^ (red) embryos (n=4-5). (**D**) Relative abundance of D- and L- 2HG (%) in the amniotic fluid samples of *Slc25a1*^-/-^ embryos compared to wild-type littermates (n=2-3). (**E**) Urine D- and L- 2HG concentration in patient samples. Data were extrapolated based on enrichment detected in two different studies. Patients 13-20 were from ref. 8; Patients 1 and 2 from ref. 11. (**F**) D-2HG enrichment in A549 cells harboring the shRNA mediated *SLC25A1* knock-down relative to control. (**G**) D-2HG enrichment in cells harboring the SLC25A1-KD or expressing IDH1**^R132C^**. (**H-I**) mRNA (**H**) and protein (**I**) levels of SLC25A1, IDH1 and IDH2 in the indicated MEFs or brain samples. (**J**) D-2HG enrichment in A549 cells harboring the SLC25A1-shRNA treated with AG120. Results were obtained from 3 experiments combined. (**K**) Expression levels of p53 and p21 in human fibroblast treated with the indicated concentrations of L- or D- 2HG. (**L**) Representative images of SA-β*-*GAL staining of control human fibroblast cells treated with indicated L-or D- 2HG for 72 hours (Bar= 100 µM). (**M-O**) Quantification of SA-β*-*GAL staining of MEFs cells (**M,O**) or control human fibroblasts (HF, **N**) treated with indicated concentrations of L-or D-2HG for 3 to 5 days (Bar= 100 µM). Bars are SD from 2 experiments. (**P**) mRNA levels of the IOS (brown) and MiDAS (green) SASPs in cells treated with D-2HG. Data are presented as mean value ± SD and p-values were calculated using unpaired t-test.

It is now emerging that wild-type IDH1/2 proteins can produce D-2HG, albeit at much lower rates and concentrations (Ježek et al, 2020). First, we studied the IDH1 and IDH2 mRNA and proteins levels and we determined that the expression of IDH1, but not of IDH2, was increased in the *Slc25a1^-/-^* MEFs and brains (Fig. 8H,I and EV3BC). Second, we employed inhibitors for IDH1 and 2 and studied their effects on D-2HG accumulation. AG120 was initially developed as an inhibitor for mutant IDH1, but more recently it was shown to have inhibitory activity at low concentrations against wild-type IDH1 in many cell lines (Zarei et al, 2022). As seen in Figure 8J, the accumulation of D-2HG was nearly entirely suppressed by AG120, but not by the IDH2 inhibitor AGI-6780 (EV3D). Similarly to AG120, the IDH1 shRNA led to a reduction of the D-2HG enantiomer in the patient derived cells, which also over-expressed IDH1 (EV3E).

In the next set of experiments we directly explored the idea that low levels of 2HG can induce p53-mediated OIS. To this end, we treated MEFs or primary human fibroblasts (HF) with 2HG. First, either D- or L-2HG induced p53 and p21 (Fig. 8K). In addition, very low doses of either enantiomer (nM or uM) induced SAβ*-*GAL (Fig. 8L-N). This effect did not increase when the concentrations were titrated up to 1- to-10 mM (Fig. 8O), reflective of tumor concentrations, thus suggesting that induction of senescence is a mechanism by which normal cells respond to low levels of 2HG. Further, treatment with D-2HG induced pro-inflammatory SASPs typical IOS, but not of MiDAS, particularly IL-1β and IL-6 (Fig.8P).

### Treatment of zebrafish embryos with 2HG partially recapitulates the phenotypes of SLC25A1 loss

The effects of 2HG on embryonic development have only been studied directly in the two-cells to blastocyst transition in murine embryos (Zhao et al, 2021). Given the predominant abundance of D-2HG in D/L-2HGA sustained by SLC25A1 dysfunction, we first sought to explore directly its contribution to embryonic development by using the zebrafish model as an experimental platform. By performing dose-response experiments, we observed no overt toxicity at the concentrations used, and we detected relevant phenotypes at concentrations ranging between 20-100 μM. The main organ affected was the brain, which was small and bulging (Fig. 9A,B and quantified in 9C). Alterations in the vertebral column and in the body curvature were also significant. In addition, many treated embryos exhibited alterations in the size of the head, jaw and eyes, larger hearts as well as pericardial edema. At an 80 μM D-2HG dose, embryos exhibited prominent disorganization of the brain tissue (Fig. 9B). Given the concomitant presence of both D/L-2HG enantiomers we also explored the effects of L-2HG alone or in combination with D-2HG, in this case at D/L ratio similar to those observed previously and typical of D/L-2HGA. Specifically, low concentrations of L-2HG alone induced a mild phenotype, which however became very severe at higher doses (Fig. 9D). The D-2HG in combination with L-2HG induced a prominent cardiovascular phenotype consisting of heart enlargement and reduced ability of the heart to pump blood (Fig. 9E and Supplemental Video 1). Further, mixing D- and L at doses at which neither enantiomer alone induced severe alterations, significantly worsened the phenotype (Fig. 9F), suggesting that the presence of both enantiomers in SLC25A1-mediated D/L-2HGDH may worsen the phenotypical alterations. These phenotypes were less prominent relatively to complete depletion of *slc25a1* with the targeted morpholino, which resulted in relatively high percent of lethality (Catalina-Rodriguez et al, 2012). We conclude that 2HG at least partially contributes to the embryonic developmental alterations seen in D/L-2HGA sustained by SLC25A1 deficiency.

**Figure 9.**
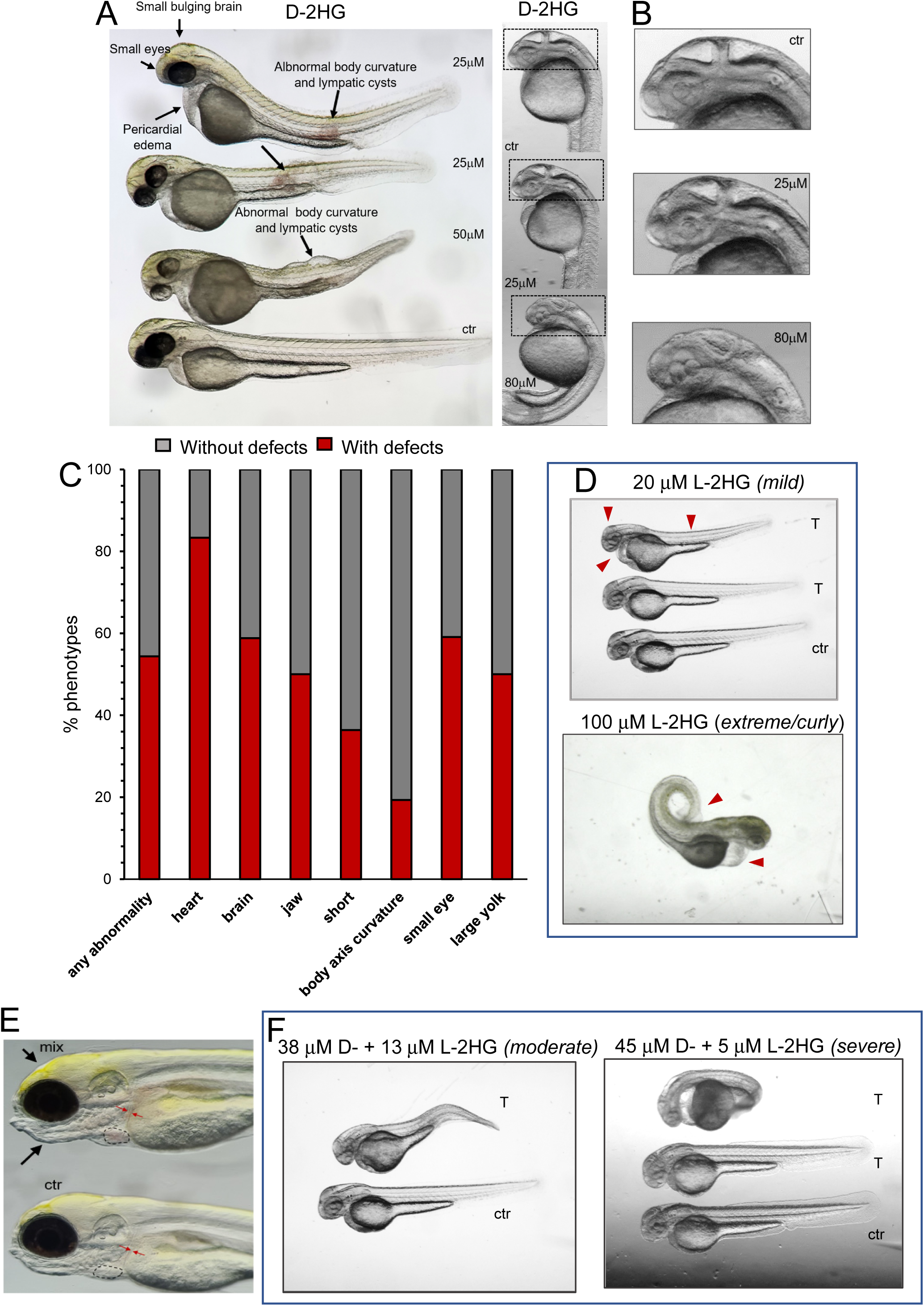
2HG perturbs embryonic development in zebrafish embryos. (**A**) Phenotypes of 25 µM or 50 µM D-2HG treated embryos at 2 dpf. Embryos were injected at blastula-stage. Prominent alterations are small eyes, bulging brains, enlarged yolk sack, pericardial edema, altered body curvature, reduced body length, lymphatic cysts, and disorganized cellular structure of tissues, most prominently seen in the brain. (**B**) Phenotypes of 25 µM or 80 µM D-2HG treated embryos at 36 hpf. Top embryo is control. Higher magnification images of the head show the dose-dependent tissue disorganization of the brain. (**C**) Percent of zebrafish with or without defects untreated or treated with 2HG (control n=4-15, 2HG treated n=18-57). Data represent the results of at least 3 repetition experiments of embryos treated with different doses of 2HG. (**D**) Phenotype of embryos treated with the indicated doses of Octyl-S-2HG at the indicated concentrations. Red arrows point to alterations in brain size, pericardial edema, enhanced body curvature and enlarged yolk. (**E**) Five days dpf Casper larvae. The top fish was treated with 30 µM D-2HG plus 20 µM L-2HG at 1 dpf. The lower fish is control. The treated larva has an enlarged heart (outlined with dashed line) and large diameter blood vessels (indicated with arrows). The treated larva additionally has a dysmorphic jaw and slightly bulging brain. (**F**) Treatment of embryos at 36 hpf, with the indicated mixtures using different ratios of D-2HG and L-2HG.

### IOS and MiDAS co-operate to restrict growth in a p53-dependent fashion

The qualitative analysis of the SASP in the brains and in the *Slc25a1^-/-^*MEFs, showed a mixed pattern of senescence involving pro-inflammatory signals derived from IOS, as well as SASPs typical of MiDAS. We have partially dissected these events as we have shown that 2HG is, *per se*, able to induce the inflammatory pro-oncogenic SASP arm, while enhanced glycolysis and decreased NAD^+^ drive mitochondrial dysfunction. In an attempt to reverse these senescence cues, we overexpressed the cDNA encoding for the D-2HGDH clearing enzyme, D-2HGDH (see EV4A for the expression levels of the protein). Overexpression of D-2HGDH reduced senescence in *Slc25a1^-/-^* MEFs (Fig. 10A). Furthermore, treating *Slc25a1^-/-^*MEFs with either NMN (Fig. 10B) or with DCA (not shown), also reduced the proportion of senescent cells. To determine the impact of these senescent programs on proliferation, we performed colony forming assays in cells harboring the SLC25A1 knock-down. While treatment with NMN or over-expression of D-2HGDH alone modestly influenced colony forming ability, the combination had a more significant additive effect enhancing colony size and number (Fig.10C and not shown). Importantly, this cooperative effect was specific for cells lacking SLC25A1 protein, as neither NMN or D2HGDH over-expression enhanced proliferation rates in the absence of the SLC25A1-KD (EVBC). Hence, IOS and MiDAS act together to restrict growth in SLC25A1 deficient cells.

**Figure 10.**
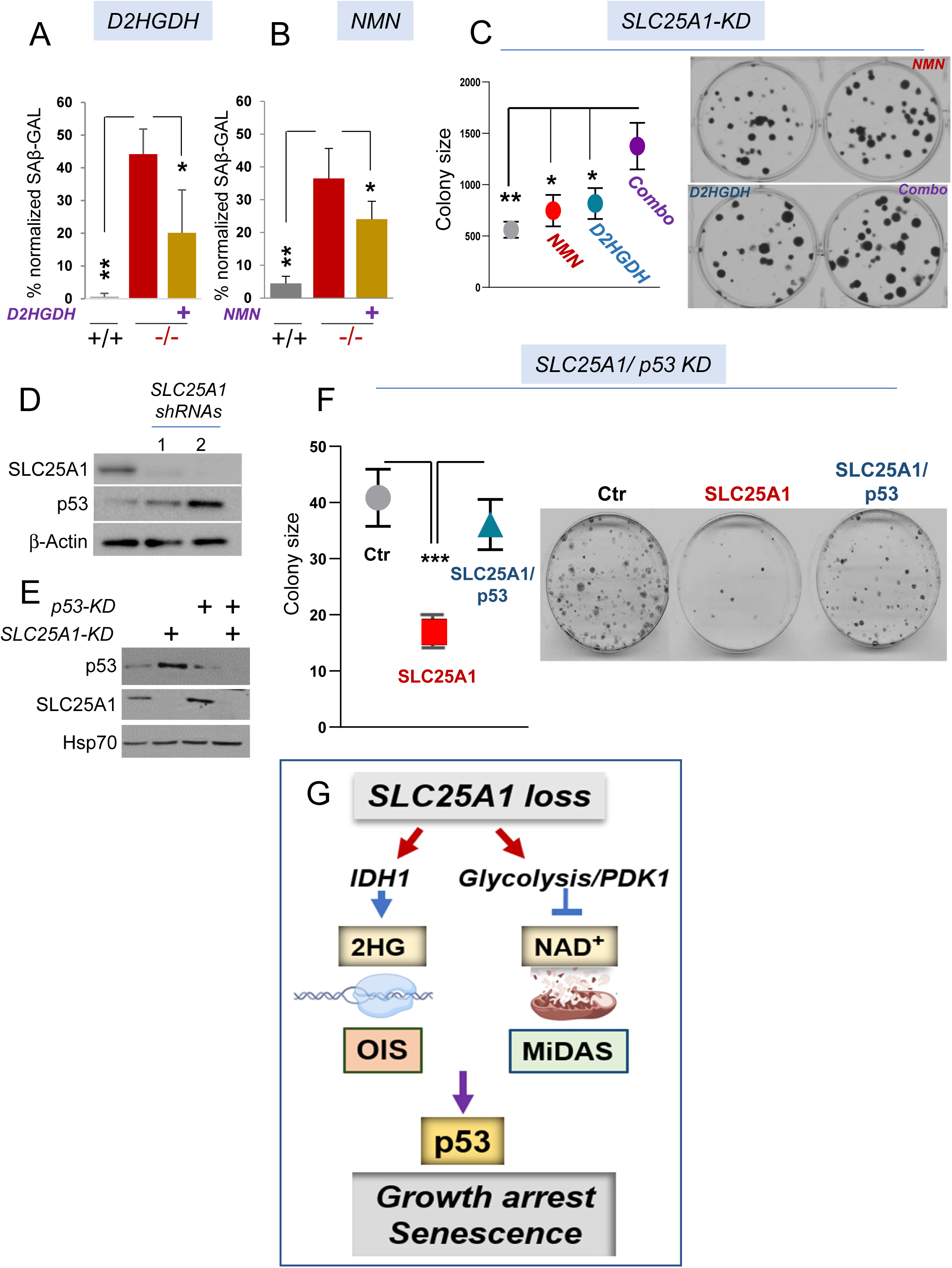
The growth defect imposed by *SLC25A1* deficiency is p53-dependent. (**A-B**) Quantification of the SAβ*-*GAL positive (%) MEF cells in the presence or absence of D2HGDH (**A**) or treated with NMN (**B**) and normalized to the total number of cells identified with DAPI counter-staining. (**C**) Quantification (with ImageJ) and representative images of colony formation assay of cells transduced with the lentivirus harboring the SLC25A1-shRNA (SLC25A1-KD) and treated with NMN, or over-expressing D2HGDH, alone or in combination (indicated as *combo*). (**D**) SLC25A1 and p53 expression levels in A549 cells transduced with the control lentivirus (pLKO, ctr) or with lentivirus vectors harboring two different SLC25A1 shRNAs (1 and 2). (**E**) Levels of expression of SLC25A1 and p53 in the double knock-down experiments performed in A549 cells. (**F**) Colony formation assay in A549 cells in the presence and absence of the Slc25a1-KD and of the p53-KD alone, or in combination. Quantification of colonies size was performed with ImageJ using multiple plates and bars represent SD. Unpaired non parametric t test was used. (**G**) Model representing the effects of *SLC25A1* loss (see also the discussion), and depicting the main pathways identified in this study leading to the induction of the IOS and MiDAS programs.

The OIS program serves as a fundamental tumor-suppressive mechanism, preventing cells that have acquired proliferative capacity from progressing through the cell cycle *via* the activation of tumor suppressors, including p53. However, cells eventually escape the senescent block by inactivating p53 and acquire the capapility to expand. First, we confirmed that the acute knockdown of SLC25A1 using two distinct shRNAs enhanced p53 expression (Fig.10D). This result demonstrates that SLC25A1 depletion directly induces p53 and raises the question of whether such induction is responsible for the proliferation defect of SLC25A1-deficient cells. To explore this concept, we employed a controlled isogenic system, A549 cells, where the effects of the SLC25A1-KD were studied in the presence or absence of the p53-KD again using colony forming assays. As shown in Figure 10E, we were able to effectively achieve a double knock-down of both proteins. The results of these experiments clearly demonstrated that the proliferation defect due to SLC25A1 depletion could be alleviated by simultaneously knocking down p53, hence reinforcing the idea that elimination of p53 activity ameliorates the proliferation defect imposed by dysfunctional SLC25A1 (Fig. 10F).

## DISCUSSION

The pathogenesis of genetically inherited human diseases due to SLC25A1 loss is still largely unclear. Using a multifaceted approach which includes a novel murine model of SLC25A1 deficiency our results provide evidence that its inactivation leads to the engagement of two pathways of senescence, OIS and MiDAS. They further show that accumulation of low doses 2HG and mitochondrial dysfunction driven by enhanced glycolysis and depletion of NAD^+^, respectively, trigger these senescent programs leading to activation of the p53 (summarized in Fig. 10G). We have also demonstated that the molecular and phenotypic alterations arising from SLC25A1 deficiency are in part sustained by the accumulation of D-2HG based on the observation that in zebrafish, treatment with this enantiomer was able to recapitulate some salient phenotypic features that overlap with the *slc25a1* depletion induced by a specific morpholino. The main impact of these findings is in the observation that SLC25A1 loss leads to only a modest enrichment of 2HG, yet these low concentrations appear to be fully pathogenic. In addition, wild-type IDH1 is responsible for the synthesis of the D-enantiomer at least in the model systems that we have studied. Finally, D-2HG induces senescence, a finding only in apparent contrast with its definition of as “oncometabolite”. Indeed, our data suggest that the first time that the response of normal cells to low doses of 2HG is the induction of senescence and that dismantling of downstream tumor suppressor pathways, including p53, is necessary for transformation, similarly to oncogene induced senescence.

The above observations have several important implications. First, consistent with the RNAseq data showing an enrichment of pro-oncogenic pathways, an important question is whether SLC25A1 loss is eventually oncogenic. It is relevant to note that as in the case of VCFS and DGS, there is a link between 2HG-acidurias and cancer risk. In fact, children affected by 2HG-acidurias sustained by mutations of *IDH1/2* or *D/L-2HGDH*, exhibit high cancer incidence, particularly of oligodendriomas, low grade gliomas, anaplastic astrocytomas and medulloblastomas (Murphey et al, 2022; Dilber et al, 2022; Srinivasan et al, 2020; Patay et al, 2012; Toer et al, 2017; Patay et al, 2015). The increased cancer risk in D/L-2HGDA sustained by SLC25A1 dysfunction may be obscured by the early death observed in these children, but deletions/mutations of the *SLC25A1* gene can be detected in various tumors (unpublished observations). Second, it is well known that the presence of proto-oncogenes in normal cells induces the activity of the p53 tumor suppressor to elicit senescence as a mechanism to halt the proliferative capacity of cells that have acquired oncogenic potential. Our data now reveal that 2HG behaves as an oncogene, inducing p53/p21-mediated OIS senescence and thus, implying that p53 interacts with potentially oncogenic metabolic products similarly to its interaction with oncogenic proteins. This is a novel concept that will offer a paradigmatic example of how alterations in the metabolism may lead to senescence and cancer. Third, our data have implications beyond diseases linked to SLC25A1 deficiency. Low levels of enrichment of 2HG are currently considered non-pathogenic, but this moiety is present in modest amount in metabolic disorders and in tumors that do not harbor IDH1/2 mutations (Du and Hu, 2021). Our results challenge this concept, revealing multifaceted mechanisms of action of 2HG depending on the cellular context, and as such, will likely drive further investigations into the broader pathogenic role of this metabolite as well as of wild-type *IDH1/2* genes in human diseases.

We have demonstrated that the IOS and MiDAS pathways are independently induced in cells lacking SLC25A1 by discernible signals. Specifically, our previously published work had demonstrated that this protein plays a fundamental role in maintaining the rates of mitochondrial respiration, as argued not only based on the deletrious effects on OXPHOS of specific inhibitors for this protein (Fernandez et al, 2020), but also given that over-expression of SLC25A1 enhances the rates of OXPHOS (Palmieri et al, 2020; Jiang et al, 2017). The finding that SLC25A1 over-expression, *per se*, drives mitochondrial oxidative metabolism, together with results presented herein, demonstrates that this activity of SLC25A1 is independent of 2HG accumulation. The mechanisms by which SLC25A1 loss leads to deterioration of OXPHOS were up to this point unclear, but we have now identified the depletion of NAD^+^ and high glycolytic rates, presumably due to the HIF1α-mediated pseudohypoxia response, as at least in part responsible for this effect in the cellular models relevant to human diseases that we have examined. It is also highly significant to note that the NAD^+^-driven pseudohypoxia response has been linked to aging (Gomes et al, 2013). Further, in the seminal work of the Campisi’s laboratory MiDAS was detected in murine models of progeroid syndromes (Wiley et al, 2016). Indeed *Slc25a1* homozygous embryos exhibit growth retardation, wrinkled skin, and facial deformities including anophtalmia, which are seen in models of progeria (Carrero et al, 2016). In addition, deletion of the gene encoding for longevity associated deacetylase, *SIRT1* leads to alterations similar to those seen in *Slc25a1^-/-^* mice, including the perinatal lethality, exencephaly, eye abnormalities, small body size and the induction of p53-mediated senescence (Liu et a, 2012). Interestingly, deficiencies of other mitochondrial transporters of the SLC25 family are known to lead to mitochondrial dysfunction, early aging and progeroid features (Rodríguez-García et al, 2018; Writzl et al, 2017). Viewed together with our findings, these observations raise the attractive possibility that the MiDAS response in the context of SLC25A1 deficiency leads to an accelerated aging, progeroid phenotype. The identification of drugs (DCA, NMN) able to reverse this deficit provides novel means of therapeutic interventions.

In summary, data presented in this work reveal that complete elimination of SLC25A1-mediated signaling during embryonic development leads to molecular and phenotypic alterations more complex than anticipated and that disorders sustained by its loss should be reclassified as dismetabolic and mitochondrial disorders. It is further relevant to note that the IOS and MiDAS pathways of senescence have been considered distinct because of the nature of their up-stream regulatory signals and their impact upon very different biological processes. Here, we identify SLC25A1 deficiency as a patho-physiological context where these signals operate together. Lastly, we have observed an enhancement of citrate-dependent pathways and lipid build-up in *Slc25a1* deficient embryos. Citrate and lipids are well known mediators of inflammation and of senescence (Hong et al, 2023), raising the question of whether a maladaptive compensatory response to replenish the cytosolic pool of citrate and lipids also contribute to the loss of proliferative capacity. These conclusions notewistanding, it is relevant to consider that the spectrum of molecular and biochemical alterations seen in *Slc25a1^-/-^* mice is global and various tissues might respond or adapt to SLC25A1 loss differently. Thus, unraveling the complexity of tissue-specific functions of SLC25A1 will be an object of future investigations. Though many of the pathological manifestations due to SLC25A1 deficiency are likely irreversible, as they are acquired during embryogenesis and development, we predict that this study will inform on therapeutic strategies able to ameliorate at least some aspects of these devastating disorders.

## Methods

### Study approval

All animal studies were conducted in compliance with ethical regulations according to protocol #2017-1192 approved by the Institutional Animal Care and Use Committee (IACUC) at Georgetown University. All the mice were housed at Georgetown University Division of Comparative Medicine, in a SPF vivarium which is maintained at a 12:12 h light:dark cycle, at 68–74°F and 30– 70% humidity range. Mice were euthanized according to the IACUC guidelines and approved protocols.

### Cell culture

The A549 and H1299 cell line were obtained from the tissue culture core facility at LCCC. Cells were grown in Dulbecco’s Modified Eagle’s Medium (DMEM 4500 mg/L glucose) supplemented with 10% fetal bovine serum, 4 mM L-glutamine, 1 mM sodium pyruvate and 1% antibiotic solution. The S*LC25A1*-specific shRNA vectors were purchased from Sigma (TRCN0000232825; TRCN0000255350) and validated before (6,18,24). Octyl-2-HG (Cayman 16366, 16367) was diluted in PBS pH 7.2.

### Mice

The S*lc25a1* heterozygous mice were purchased from the Mutant Mouse Resource Research Center (MMRRC) (C57BL/6N-*Slc25a1^tm1a(EUCOMM)Wtsi^*, RRID:MMRRC_042258-UCD). The targeting vector allows for constitutive or conditional deletion of the *SLC25A1* gene through the incorporation of an *IRES:lacZ* trapping cassette which is inserted between introns 1 and 5 of the *Slc25a1* gene [24]. Mice were maintained on a 12h light/dark cycle and were supplied with food and water ad libitum. To obtain mouse lacking *Slc25a1* we crossed heterozygous *Slc25a1*^+/−^ heterozygous mice. Pregnancy was confirmed by checking for vaginal plugs and the increase in body weight measured weekly.

### Embryos, brains and amniotic fluids collection

Pregnant mice were euthanized at different stages of pregnancy and whole embryos were collected. Typically, embryos were harvested at E9.5-E19, and tips of tails were collected for genotyping. For IHC, embryos were fixed in 4% paraformaldehyde overnight, transferred to 80% EtOH and sent for histological processing. Paraffin embedding, sectioning and hematoxylin/eosin staining of embryos tissue sections were performed by Histology and Microscopy Core of Georgetown University or by VitroVivo Biotech. To collect the amniotic fluid, and avoid cross-contamination, we typically selected embryos distally located. After transferring the embryos to a clean dish, the amniotic fluid was collected by 1-mL syringe and 27-gauge needle. For biochemical and molecular analysis, indicated organs were dissected, quickly washed in PBS and snap frozen in liquid nitrogen. For the collection of the brains, skin and skull (when present) were carefully removed. The isolated brains were quickly washed in PBS twice, snap frozen in liquid nitrogen and stored at -80°C until processed for further analyses.

### CT Images acquisition from the International Mouse Phenotyping Consortium

The IMPC has provided CT images of *slc25a1^-/-^* mice without description of the phenotypes. Analysis of these images and phenotype identification was performed in house at GUMC through the help of the Preclinical Imaging Research Laboratory (PIRL) facility.

### 2HG quantification with LC/MS/MS and GS/MS/MS

For LC/MS a targeted metabolomics method was used to quantitate 2-HG using QTRAP® 7500 LC-MS/MS System (Sciex, MA, USA). The 2HG standard was from Sigma Aldrich (# 90790-10MG). For the purpose, 75 μL of extraction buffer (methanol/water 50/50) containing 200 ng/mL of 4-nitrobenzoic acid as internal standard for negative mode was added to the cell pellet and sample tube was plunged into dry ice for 30 sec and 37 °C water bath for 90 sec. This cycle was repeated for two more times and then samples were sonicated for 1 minute. The samples were vortexed for 1 min and kept on ice for 20 minutes followed by addition of 75 μL of ACN. The samples were incubated at -20 °C for 20 minutes for protein precipitation. The samples were centrifuged at 13,000 rpm for 20 minutes at 4 °C. The supernatant was transferred to MS vial for LC-MS analysis. 20 μL of each prepared sample was mixed to generate the pooled QC sample. For GC-TOF amniotic fluids or tissue samples were processed using 500 μL of 50% methanol in water containing the internal standard (4-Nitrobenzoic acid, prepared in MeOH at a concentration of 20 µg/mL). The samples were homogenized on ice to ensure tissue lysis and metabolite extraction was then vortexed for 2 minutes. A volume of 10 µL was withdrawn from the homogenized tissue suspension for the protein quantification assays. The samples were vortexed for 2 minutes and left for 2 hours at room temperature. The BCA kit used for the protein quantification assay was Pierce™ BCA Protein Assay Kit (Cat # 23225). A volume of 20 μL of methoxyamine (20 mg/mL) was added to the dried tissue extracts and then heated in an agitator at 60 ^0^C for 30 minutes. This was followed by 100 μL of MSTFA. The vials were transferred to an agitator to heat at 60 ^0^C for 30 more minutes. Finally, the vials were capped and a volume of 1.5 μL was injected directly to the GC-MS. The samples were allowed to react at room temperature for 20 minutes before being transferred to the GC for injection. Briefly after the derivatization process, a volume of 1.5 µL of the derivatized solution was injected in (1:10) split mode into an Agilent 7890B GC system (Santa Clara, CA, USA) that was coupled with a Pegasus HT TOF-MS (LECO Corporation, St. Joseph, MI, USA). Separation was achieved on a Rtx-5 w/Integra-Guard capillary column (30 m x 0.25 mm ID, 0.25 μm film thickness; Restek Corporation, Bellefonte, PA, USA), with helium as the carrier gas at a constant flow rate of 0.9 mL/min. The temperature of injection, transfer interface, and ion source were set to 150, 270, and 320 °C, respectively. The GC temperature programming was set to 0.2 minutes of isothermal heating at 70 °C, followed by 6 °C/min oven temperature ramping to 270 °C, a 7.0 minute isothermal heating of 270 °C, 20 °C/min to 320 °C, and a 2.0 min. isothermal heating of 320 °C. Electron impact ionization (70 eV) at full scan mode (40–600 *m*/*z*) was used, with an acquisition rate of 20 spectra per second in the TOF/MS setting.

### Metabolomic and Lipidomics

Metabolomic (targeted and untargeted was performed as described before (Tan et al, 2020). The lipidomic method used in the brains of *Slc25a1^-/-^* mice, is designed to measure 19 classes of lipid molecules which includes diacylglycerols (DAG), chloesterol esters (CE), sphingomyelins (SM), phosphtatidylchloine (PC), triacylglycerols (TAG), free fatty acids (FFA), ceramides (CE), dihydroceramides (DCER), hexosylceramide (HCER), lactosylceramide (LCER), phosphatidylethanolamine (PE), lysophosphtatidylchloine (LPC), lysophosphatidylethnolamine (LPE),phosphatidic acid (PA), lysophosphatidic acid (LPA), phosphatidylinositol (PI), lysophosphotidylinositol (LPI), phosphatidylglycerol (PG) and phosphatidylserine (PS) using QTRAP® 5500 LC-MS/MS System (Sciex). For the purpose, 20 µL of each amniotic fluid sample was dissolved in 100 μL of chilled isopropanol containing internal standards was added and samples were vortexed. The samples were vortexed again for 1 min and kept on ice for 30 minutes. Samples were incubated at -20 °C for 2 hours for complete protein precipitation. The samples were centrifuged at 13,000 rpm for 20 minutes at 4 °C. The supernatant was transferred to MS vial for LC-MS analysis. Five µL of each sample was injected onto a Xbridge amide 3.5 µm, 4.6 X 100 mm (waters) using SIL-30 AC auto sampler (Shimazdu) connected with a high flow LC-30AD solvent delivery unit (Shimazdu) and CBM-20A communication bus module (Shimazdu) online with QTRAP 5500 (Sciex, MA, USA) operating in positive and negative ion mode. Obtained results were analyzed using MetaboAnalyst.

### Brain TCA cycle analyses

Tissue samples were mixed with 400 µL of MeOH containing the internal standard (4-Nitrobenzoic acid) which was prepared in MeOH at a concentration of 30 µg/mL and homogenized for 60 seconds. A volume of 10 µL was withdrawn from the tissue suspension for the protein concentration assay. The samples were mixed for 2 minutes, left at -20 °C. for 2 hours then centrifuged for 30 min at a speed of 16,000 X g and a temperature of 4 °C. The supernatant was separated and transferred to the GC vials then dried under vacuum at room temperature. Samples were derivatized and injected into GCMS instrument. Protein concentration was determined using BCA assay. The Briefly after the derivatization process, a volume of 2.0 µL of the derivatized solution was injected in a splitless mode into an Agilent 7890B GC system (Santa Clara, CA, USA) that was coupled with a Pegasus HT TOF-MS (LECO Corporation, St. Joseph, MI, USA). Separation was achieved on a Rtx-5 w/Integra-Guard capillary column (30 m x 0.25 mm ID, 0.25 μm film thickness; Restek Corporation, Bellefonte, PA, USA), with helium as the carrier gas at a constant flow rate of 0.8 mL/min. The temperature of injection, transfer interface, and ion source was set to 150, 270, and 320 °C, respectively. The GC temperature programming was set to 0.2 min. of isothermal heating at 70 °C, followed by 6 °C/min oven temperature ramping to 300 °C, a 1.0 min. isothermal heating of 300 °C, 20 ° C/min to 320 °C, and a 2.0 min. isothermal heating of 320 °C. Electron impact ionization (70 eV) at full scan mode (40– 600 m/z) was used, with an acquisition rate of 20 spectra per second in the TOF/MS setting.

### Genotyping

Mouse tail biopsies and agarose gel electrophoresis was used for analysis of PCR products. The PCR conditions used were as follows: 94 °C for 1 minute, 5 cycles at 94 °C for 30 s, 63 °C for 40s, 68 °C for 1 min; followed by 30 cycles (94 °C for 30 seconds, 55 °C for 40 seconds, and 72 °C for 1 min), 72 °C for 7 min then kept in 4 °C before use. The primers used for genotyping are indicated in EV Table 1.

### Mouse embryonic fibroblasts isolation

Freshly dissected embryos at E10-E14 were used for isolation of mouse embryonic fibroblasts (MEFs). Brain, liver, internal organs and tail were removed, and the remaining tissues were rinsed with PBS and transferred to a 10 cm plate. Embryos were then minced with a sterile razor blade in the presence of TrypLE™ Express Enzyme (Gibco™, 12604021) following by 10 min incubation on a shaker at RT. Finally, the cells were pipetted multiple times to further disintegrate the tissue, and then the freshly isolated cell were cultured in high glucose DMEM medium (Dulbecco’s Modified Eagle Medium with 10% fetal bovine serum, 4500 mg/L glucose, 4 mM L-glutamine, 1 mM sodium pyruvate and 1% antibiotic solution).

### Protein analysis and immunoblot

Proteins from cells or frozen tissue samples, grinded in liquid nitrogen, were homogenized in RIPA buffer supplemented with cOmplete Mini Protease Inhibitor Tablets (Roche, 11836153001). Protein quantification was done by using Coomassie (Bradford) Protein Assay Kit (Pierce). Equal amount of protein lysate was loaded and separated in an SDS-polyacrylamide gel electrophoresis using Novex™ 4-20% Tris-Glycine Mini Gel (Invitrogen, XP04200PK2) and transferred to PVDF membrane (Millipore, IPVH304F0). The membrane was blocked in blocking buffer with 10% horse serum to prevent non-specific binding, followed by incubation with different antibodies: SLC25A1 (Santa Cruz, sc-86392 or Proteintech, 15235-1-AP), ACC1 (Cell Signaling, #3676), FASN (Cell Signaling, #3180), IDH1 (Proteintech, 12332-1-AP), AMPK-α (Cell Signaling, #5832), pAMPK-α (Cell Signaling, #2535), Chk1 (Cell Signaling, Cat#2360), HIF1-alpha (Novus Biological, Cat#NB100-105), p21 Waf1/Cip1 (Cell Signaling, #37543), mTOR (Cell Signaling, #2983), pH2A.X (Cell Signaling, #9718), p53 (Santa Cruz, sc-126 and Sigma Aldrich, OP29-100UG), D2HGDH (Proteintech, 13895-1-AP). Appropriate horseradish peroxidase–conjugated secondary antibody (Invitrogen) was applied after incubation with primary antibody, SuperSignal™ West Pico PLUS Chemiluminescent Substrate (Thermo Scientific, 34577) or SuperSignal™ West Dura Extended Duration Substrate (Thermo Scientific, 34075) were used for protein detection. β-Actin (Santa Cruz, sc-47778) and HSP70 (Santa Cruz, sc-24) served as a loading control.

### Quantitative real-time PCR

Total RNA was extracted from frozen tissue and cells pellet samples using TRIzol™ Reagent (Invitrogen, 15596026), followed by RNA quantification with NanoDrop (Implen, Munich, Germany). After genomic DNA treatment by DNase I (Invitrogen, AM2222), total RNA was used in cDNA synthesis with SuperScript™ IV Reverse Transcriptase (Invitrogen, 18090200) and random hexamers according to the manufacturer’s instructions. The quantitative Reverse Transcriptase-PCR (qRT-PCR) was performed on the QuantStudio™ 12K Flex Real-Time PCR System (Applied Biosystems) with PowerUp™ SYBR Green Master Mix (Applied Biosystems). The obtained results were normalized to housekeeping genes, and their sequences will be provided.

### RNA sequencing

Brains isolated from 18.5 dpf embryos were used for RNA sequencing. Briefly, RNA isolatation and RNA sequencing (RNAseq) was performed by Novogene. Raw intensity data was background corrected and filtered of low expression genes across samples (low expression genes across samples (low expression genes across samples (<100 read) prior to analysis. Normalization and differential expression were performed in R using the DESeq2 package. A cutoff of p < 0.05 and |FC| > 2 were used to determine differentially expressed genes. The GSEA Java-based software package from The Broad Institute was used to identify top enriched pathways in *Slc25a1*^+/+^ and *Slc25a1*^-/-^ brains. Reference genome and gene model annotation files were downloaded from genome website directly. Index of the reference genome was built using Hisat2 v2.0.5 and paired-end clean reads were aligned to the reference genome using Hisat2 v2.0.5. featureCounts v1.5.0-p3 was used to count the reads numbers mapped to each gene. FPKM of each gene was calculated based on the length of the gene and reads countmapped to this gene. FPKM, expected number of Fragments Per Kilobase of transcript sequence per Millions base pairs sequenced, considers the effect of sequencing depth and gene length for the reads count at the same time, and is currently the most commonly used method for estimating gene expression levels. Differential expression analysis was performed using the DESeq2 R package (1.20.0). The P values were adjusted using the Benjamini & Hochberg method. Corrected P-value of 0.05 and absolute fold change of 2 were set as the threshold for significantly differential expression. Enrichment analysis of differentially expressed genes.

### Immunohistochemistry and immunofluorescence

Age-matched embryos were removed from the uteri and fixed in 4% paraformaldehyde in phosphate-buffered saline and paraffin embedded, sectioned in the mid sagittal plane and stained with hematoxylin and eosin (H&E) and anti-Slc25a1 antibody (Proteintech, 15235-1-AP), used at 1/150 dilution. Consecutive sections with the primary antibody omitted were used as negative controls. Stained slides were scanned using the VS120 Virtual Slide Microscope (Olympus). For immunofluorescence (IF) staining, FFPE embryo sections were deparaffinized and sodium citrate was used for antigen retrieval. Slides were washed in PBS, blocked using TBST+20% Aquablock, and incubated with primary antibodies: anti-Slc25a1 (Proteintech, 15235-1-AP) and p21 (Santa Cruz, sc-756). Finally, samples were washed in PBST and incubated with appropriate secondary antibody and DAPI or Hoechst (Sigma-Aldrich). IF samples were acquired using a Zeiss LSM800 laser scanning confocal microscope and analyzed using ImageJ/Fiji software.

### Electron microscopy

Pregnant females were euthanized and perfused with 2.5% glutaraldehyde and 2% PFA in 0.15M cacodylate buffer. Following osmication, dehydration, and embedding in Epon 814, embryos heads were sectioned transversely (at the midpoint) at 120 nm using an ultramicrotome. Ultrathin sections were placed in silicon wafers and carbon taped in aluminum stubs for scanning electron microscopy (SEM) imaging in a Helios NanoLab 660 dual beam microscope. To maximize the collection of the backscattered electrons and produced TEM-like images, a high contrast solid-state backscatter electron detector, in magnetic immersion mode and 4 μm working distance, was used. The acquisition was performed using 2 kV and 0.40 nA as the current landing. Overview images including the entire section were taken at a magnification of 5,000× and analyzed using ImageJ software.

### Senescence-associated β-Gal staining

Cells grown on a 6-well plate were washed with PBS, fixed with fixative solution and stained with x-gal according to the manufacturer protocol (Cell Signaling, #9860). After counterstaining with DAPI, cells were imaged using an Olympus IX-71 Inverted Epifluorescence Scope and multiple images of each well were taken for quantification.

### Colony formation assay

H1299 and p53H1299 (or indicated cells) were plated in a six-well plate at density 500 cells/well for 14 days, and the medium was replaced every 3 days. Additionally, indicates cells were treated with nicotinamide mononucleotide (NMN, Selleckchem, catalog No.S5259) 24h after plating for 13 days, and the medium was replaced every 3 days. Cells were fixed in 4% paraformaldehyde, stained with crystal violet and imaged/photographed to visualize the results. Quantification was done by manual counting and measuring stained area with image J.

### Lentivirus production and infection

Control empty vector and vector shIDH1 were purchased from Sigma. Lentiviruses were produced in Lenti-X 293T cells with Third Generation Packaging Mix (Applied Biological Materials Inc., Cat#LV053) according to the manufactureser’s instruction. Cells were transduced with lentiviruses and Polybrene Infection/Transfection Reagent (Millipore) reagents for 24 hrs and 0.5 µg/ml puromycin was used to obtain stably expressing cell lines.

### D2HGDH overexpression (Plasmids and transfection)

D2HGDH expression plasmid (OriGene, RC207367) and pRc/CMV expression plasmid (Invitrogen) were used. Cells were plated in a six-well plates and allowed to grow overnight, and transfected using Lipofectamine 2000 (Invitrogen, cat. 11668-030) and OPTI-MEM reduced serum media (Gibco, cat. 31985-062). The medium was replaced after 24 h. The D2HGDH overexpression was verified by Western blotting using D2HGDH antibody (Proteintech, 13895-1-AP).

### Seahorse extracellular flux analyzer measurements

Cells were seeded at 10 000 cells/well density in high glucose DMEM with 10% FBS, 1 mM pyruvate and 1% antibiotic solution, and the following day treated with the indicated drug/vehicle for 24 h. Seahorse measurements were done according to the manufacturer’s instructions. Briefly: for the mitochondrial stress test, the medium was replaced with DMEM without FBS or bicarbonate, containing 5 or 10 mM glucose, 1 mM pyruvate, 2 mM glutamine and placed in a CO2 free incubator at 37 °C for 1 h, transferred to the Xf96 extracellular flux analyzer (Agilent). The program consisted of three measurements of OCR/ECAR before the injection of each drug: oligomycin (0.5 μM final concentration), FCCP (2 μM) and rotenone/antimycin (0.5 μM of each).

### ETC Respiratory Complex activities

The activities of the ETC were assessed with the Seahorse analyzer with injections to study individual complexes as follows. For complex I after mild permeabilization, injection 1: Pyruvate 10mM, Malate 5mM; injection 2: Rotenone. For complex II: injection 1: Rotenone (to inhibit complex I), injection 2: Succinate (10mM), Injection 3: Atpenin A5 (complex II inhibitor). For complex III: injection 1: pyruvate 10mM, injection 2: antimycin. For complex IV:injection 1: pyruvate 10mM injection 2: sodium Azide (200uM). ATP synthase: injection 1: pyruvate 10mM, injection 2: oligomycin.

### 2HG analysis

The α-Hydroxyglutaric Acid was obtained from Cayman Chemicals. Diacetyl-L-tartaric anhydride (DATAN) was purchased from FisherScientific. All other reagents and solvents were of analytical and LCMS grade. Sample preparation: a volume of 50 μL from amniotic fluid was extracted with 250 μL of methanol containing 0.004 mmol/L of the internal standard (IS). Cell pellets were extracted with a volume of 1 mL of methanol containing 0.004 mmol/L of the internal standard. Samples were centrifuged for 20 minutes at 14, 000 X g and 4 °C. The supernatant was separated in a clean set of vials. The vial contents were evaporated to dryness at 22 °C by a gentle flow of nitrogen. The di-acetyltartaryl derivative was prepared by treating the dry residue with 50 μL of freshly made 50 mg/mL DATAN in Acetonitrile–acetic acid (4:1 by volume). The vials were capped and heated at 75 °C for 60 minutes. After the vial were cooled to room temperature, the mixture was evaporated to dryness by a nitrogen stream at room temperature, the residue was redissolved in 100 μL of distilled water, and 10 μL of the aqueous solution was injected on the LC column and analyzed by LCMS. The sample, prepared in the first step, were analyzed on a Sciex QTRAP 4500 mass spectrometer equipped with a Shimadzu Prominence UFLC XR System whereas, it was separated Phenomenex Lux 3 µm AMP, LC Column 150 x 3.0 mm, PN: 00F-4751-Y0. The column temperature was kept at 22 0C. Solvent A was 100% LCMS grade water with 0.1% formic acid and solvent B was 100% LCMS grade acetonitrile with 0.1% formic acid. The source temperature was kept at kept at 220 0C. The ion spray voltage was -4500 volts. The flow rate was consent throughout the method and was set to 0.3 mL/min. The mass spectrometry method was set to a Negative MRM mode with a cycle time of 0.99 second. The targeted masses of the derivatized 2-HG were m/z 363, 147, and 129 Da. The Entrance Potential (EP) was set to -10 volts and the Declustering Potential (DP), the Collision Cell Exit Potential (CXP) were set to values optimized previously for each and every assigned transition. The concentration of the analyte in urine was calculated by interpolation of the observed analyte/IS peak-area ratio based on the linear regression line for the calibration curve, which was obtained by plotting peak-area ratios vs analyte concentration. Data was analyzed using Analyst 1.7 and Sciex OS software for quantification.

### D/L-2HG Derivatization method

The α-Hydroxyglutaric Acid was obtained from Cayman Chemicals. Diacetyl-L-tartaric anhydride (DATAN) was purchased from FisherScientific. All other reagents and solvents were of analytical and LCMS grade. A volume of 50 μL from amniotic fluid was extracted with 250 μL of methanol containing 0.004 mmol/L of the internal standard (IS). Cell pellets were extracted with a volume of 1 mL of methanol containing 0.004 mmol/L of the internal standard. Samples were centrifuged for 20 minutes at 14, 000 X g and 4 °C. The supernatant was separated in a clean set of vials. The vial contents were evaporated to dryness at 22 °C by a gentle flow of nitrogen. The di-acetyltartaryl derivative was prepared by treating the dry residue with 50 μL of freshly made 50 mg/mL DATAN in Acetonitrile– acetic acid (4:1 by volume). The vials were capped and heated at 75 °C for 60 minutes. After the vial were cooled to room temperature, the mixture was evaporated to dryness by a nitrogen stream at room temperature, the residue was redissolved in 100 μL of distilled water, and 10 μL of the aqueous solution was injected on the LC column and analyzed by LCMS. The sample, prepared in the first step, were analyzed on a Sciex QTRAP 4500 mass spectrometer equipped with a Shimadzu Prominence UFLC XR System whereas, it was separated Phenomenex Lux 3 µm AMP, LC Column 150 x 3.0 mm, PN: 00F-4751-Y0. The column temperature was kept at 22 0C. Solvent A was 100% LCMS grade water with 0.1% formic acid and solvent B was 100% LCMS grade acetonitrile with 0.1% formic acid. The source temperature was kept at kept at 220 0C. The ion spray voltage was -4500 volts. The flow rate was consent throughout the method and was set to 0.3 mL/min. The mass spectrometry method was set to a Negative MRM mode with a cycle time of 0.99 second. The targeted masses of the derivatized 2-HG were m/z 363, 147, and 129 Da. The Entrance Potential (EP) was set to -10 volts and the Declustering Potential (DP), the Collision Cell Exit Potential (CXP) were set to values optimized previously for each and every assigned transition. The concentration of the analyte in urine was calculated by interpolation of the observed analyte/IS peak-area ratio based on the linear regression line for the calibration curve, which was obtained by plotting peak-area ratios vs analyte concentration. Data was analyzed using Analyst 1.7 and Sciex OS software for quantification.

### Zebrafish

All zebrafish procedures were performed in accordance with NIH guidelines for the maintenance and use of laboratory animals and approved by the Georgetown University Institutional Animal Care and Use Committee Protocol #2017-0078. Mixed wild-type, or *mitfa^w2/w2^; mpv17 ^a9/a9^* (casper), embryos were obtained by pair-wise and group breeding, and raised in fish water (0.3 g/L sea salts) at 28°C. For drug treatments, (2R)-Octyl-〈-hydroxyglutarate (Cayman Chemicals) and (2S)-Octyl-〈- hydroxyglutarate (Cayman Chemicals), solutions were prepared in fish water. Embryos were treated at blastula, 1 dpf, or 2 dpf stages at a density of 100ul drug solution per embryo. Chorions were removed at 1 dpf by 10 min incubation in 200ug/ml pronase. Drug solutions were replaced daily. SA– β–GAL staining was performed as described (Da Silva-Álvarez et al., 2020) modified with the replacement of X-gal with Bluo-Gal, 5-Bromo-3-indolyl β-D-galactopyranoside.

### Statistical analyses

For the analysis of the RNAseq experiments in the brain, differentially Expressed Genes were obtained using DESeq2 (2). Significantly upregulated Genes (FDR < 0.05) were assessed for pathway enrichment using the Reactome 2022 database (5) via EnrichR (4). Significantly enriched Pathways were clustered via GeneSetCluster(3) using a distance score that reflects similarities in genes mapping to multiple significantly enriched pathways. Similar processes were summarized by secondary annotation. The p-values of Reactome pathway enrichment were displayed as a Heatmap via GraphPad Prism. For all experiments, results are presented as mean value ±SD or SEM, unless specified otherwise. Statistical significance was assessed using unpaired non parametric t-test, unless indicated otherwise. Significant differences were graphed using GraphPad Prism software and p< 0.05 was considered as statistically significant.

## Data availability

All data will be made available on public databases and upon request.

## Authors contribution

MLA designed all experiments and wrote the paper. JV accrued the patients derived cell lines from patients under his care. AKP performed the majority of the experiments, organized the data for the Figures and proof-edited the manuscript for accuracy in the description of the experiments. MT generated the mouse colonies and worked with AKP for three years on this project. RR performed the IHC and the IF experiments. AMI designed and performed the statistical analyses on the transcriptomic data. HF performed the Seahorse experiments in the *Slc25a1*^-/-^ MEFs and contributed to the initial functional characterization of the mice-derived cells. RM has maintained all the mouse strains during Covid 19, and performed several experiments. GWP provided supervision to RR with the IF experiments. EG and MLA designed the experiments in zebrafish, EG performed the experiments, assembled the Figures and provided interpretation of the results. CA provided expertise in the interpretation of the phenotype and of the CT scans of the mice, proof-edited the manuscript and offered inputs on this project throughout the years.

## Acknowledgements

Studies on SLC25A1 in the MLA lab have been supported by R01CA193698, R21DE028670 and R21CA256546. JV is the Cleveland Family Endowed Chair in Pediatric Research with support in part from NIH grant R01DK109907. All these studies have been supported by the outstanding facility at GUMC and by the 2P30CA051008-30 CCSG grant. We are grateful to Dr. Anton Wellstein for many inputs on this project, to the Riegel/Wellstein and to Dr. Chunling Yi laboratories for sharing reagents and instrumentation. We thank Drs. Michael Girgis and Amrita Cheema for the outstanding contributions for the metabolomic analyses and for the derivatization of D/L-2HG. We are also very grateful to Anastas Popratiloff and Cheryl Clarkson for the ECM analysis and to Lawrence F. Kroemer for initial aid with the assessment of the brain alterations in the mice. We thank Sarah Flowers for her inputs on all of the projects related to SCL25A1.

