## Supplemental figures and legends for "Loss of the mitochondrial carrier, *SLC25A1,* during embryogenesis induces a unique senescence program controlled by p53"

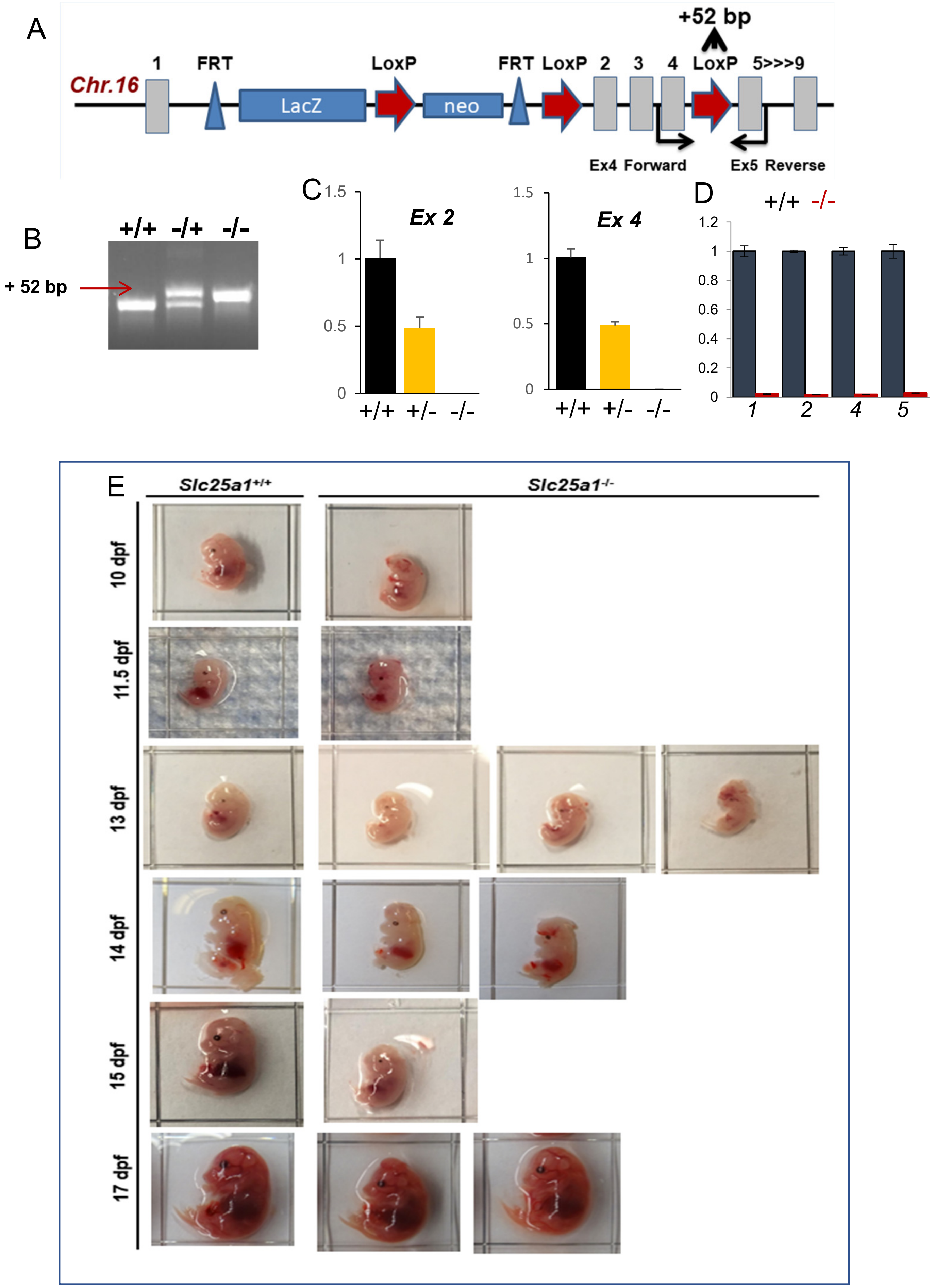

A
+52 bp
1
FRT
LoxP
FRT
LoxP
2
3
4
LoxP
5>>>9
Chr.16
LacZ
neo
Ex4 Forward
Ex5 Reverse
B
+/+
-/+
-/-
+ 52 bp
C
Ex 2
Ex 4
1.5
1
0.5
0
+/+
+/-
-/-
D
+/+
-/-
1.2
1
0.8
0.6
0.4
0.2
0
1
2
4
5
E
Slc25a1+/+
Slc25a1-/-
10 dpf
11.5 dpf
13 dpf
14 dpf
15 dpf
17 dpf

**EV1. Slc25a1 deficient embryos can be recovered at all stages of embryonic development.**

(**A-B**) Schematic representation of the knock-out first allele *tm1a* cassette targeting the *mus musculus Slc25a1* gene on chromosome 16 (**A**). The insertion of the *LoxP* site between *exons* 4 and 5 in chromosome 16 creates a difference of 52 bp that allows distinguishing of the +/+; +/-; and -/- mice (**B**).

(**C,D**). Quantitative rtPCR with primers spanning *exons 2* and *4* of the Slc25a1 transcript in the brain (**C**) and *exons 1, 2, 4, 5* in MEFs (**D**).

(**E**) Representative images of *Slc25a1*$^{+/+}$ and *Slc25a1*$^{-/-}$ embryos collected at the indicated stages of embryonic development. Identical scale in each set.

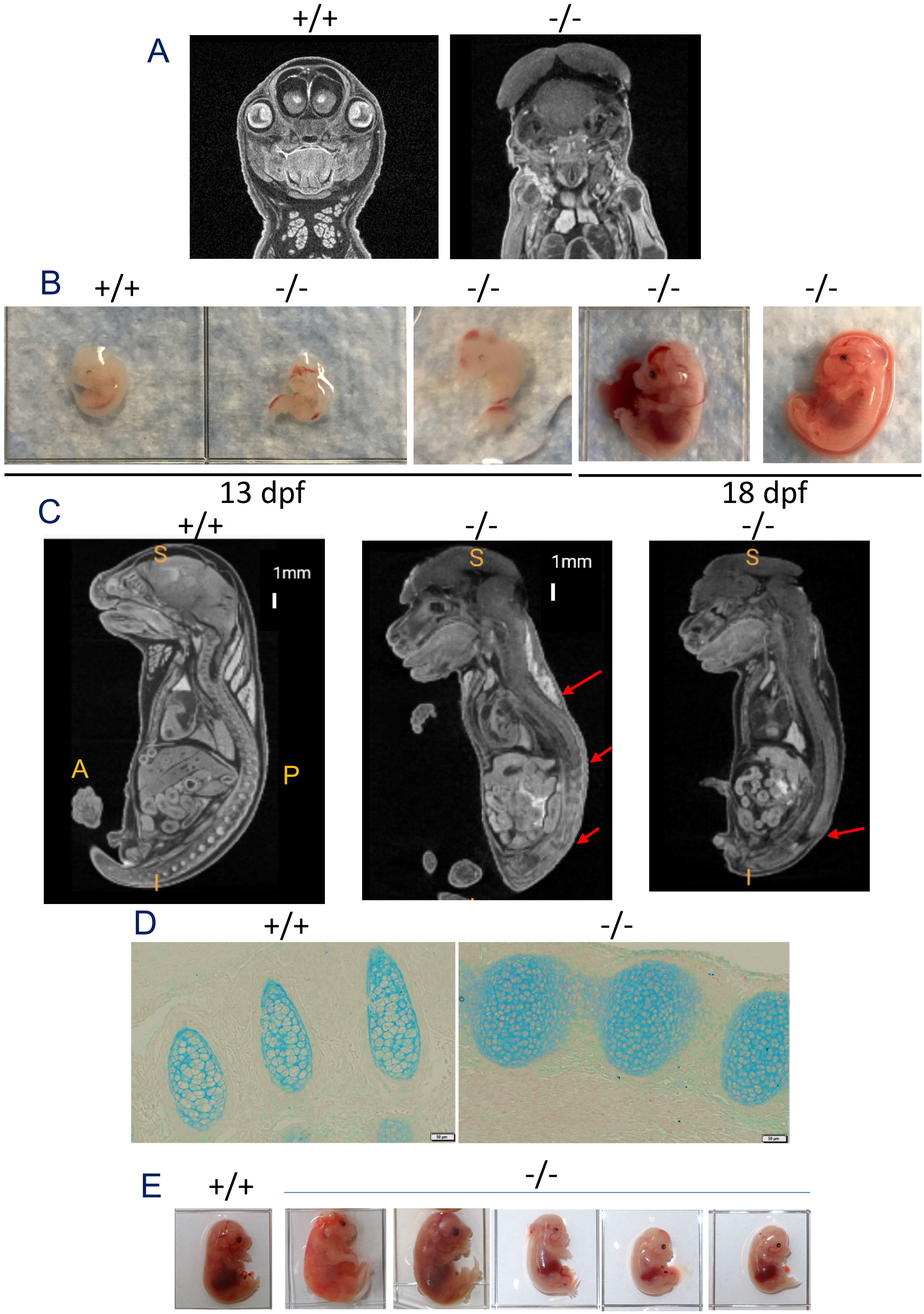

A
+/+
-/-
B
+/+
-/-
-/-
-/-
-/-
13 dpf
18 dpf
C
+/+
-/-
-/-
S
1mm
A
P
I
S
1mm
S
I
D
+/+
-/-
E
+/+
-/-

**EV2.** Heterogeneity of the phenotypes induced by *Slc25a1* loss.

**(A)** Facial dysmorphic features of *Slc25a1*$^{-/-}$ embryos at E18.5 dpf compared to wild-type and detected with CT scan (UC Davis library).

**(B)** Prominent hemorrhagic phenotype in *Slc25a1*$^{-/-}$ embryos affecting the brain, liver and abdominal organs.

(**C-D**) Alterations of the vertebral column in *Slc25a1*$^{-/-}$ embryos at E18.5 dpf, including abnormal curvature, increased thickness and poor separation of the vertebrae (**C**) and alcian blue staining of cartilage in the vertebral column (**D**).

(**E**) Phenotype of the five *Slc25a1*$^{-/-}$ embryos used for the metabolomic studies. One representative wild-type embryo is also shown.

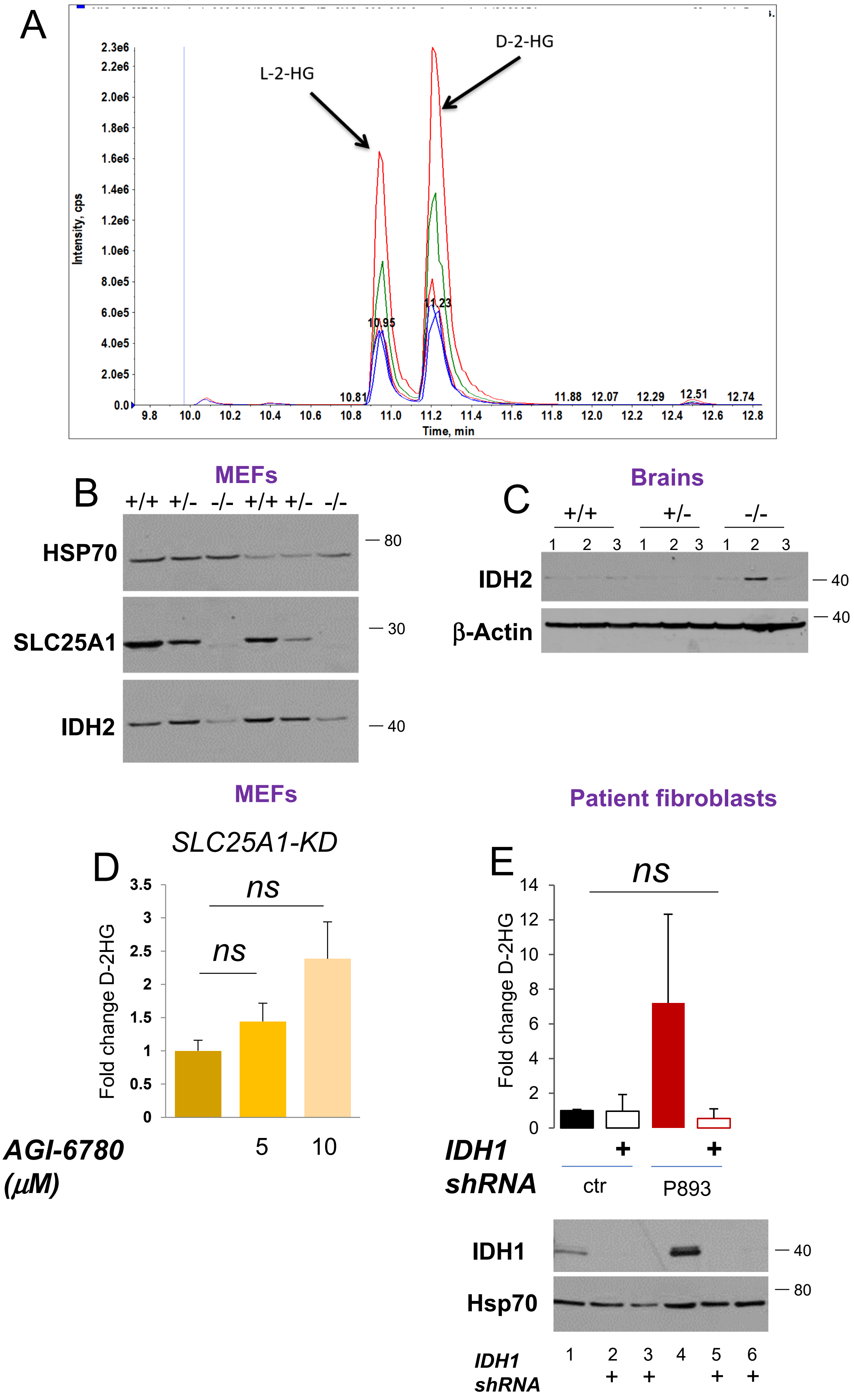

A
L-2-HG
D-2-HG
Intensity, cps
Time, min
10.81
10.95
11.23
11.88
12.07
12.29
12.51
12.74
B
MEFs
+/+ +/- -/- +/+ +/- -/-
HSP70
SLC25A1
IDH2
80
30
40
C
Brains
+/+ +/- -/-
1 2 3 1 2 3 1 2 3
IDH2
β-Actin
40
40
MEFs
SLC25A1-KD
D
ns
ns
Fold change D-2HG
AGI-6780 (μM)
5 10
Patient fibroblasts
E
ns
Fold change D-2HG
IDH1 shRNA
+ +
ctr
P893
IDH1
Hsp70
40
80
IDH1 shRNA
1 2 3 4 5 6
+ + + +

**EV3. Characterization of D-2HG in SLC25A1 deficient cells.**

(**A**) Peak separation of L- and D- 2HG using a chiral derivatization method with diacetyl-L-tartaric anhydride. The di-acetyltartaryl derivative was prepared by treating the dry residue with 50 µL of freshly made 50 mg/mL DATAN in Acetonitrile–acetic acid (4:1 by volume) and samples were analyzed on a Sciex QTRAP 4500 mass spectrometer equipped with a Shimadzu Prominence UFLC XR System ((see also materials and methods). X-axis represents the time in minutes the y-axis is the intensity in cps units (counts per second).

(**B-C**) Expression levels of IDH2 and SLC25A1 in the MEFs (**B**) and brains (**C**) of the indicated mice.

(**D**) Levels of enrichment of D-2HG in cells treated with the IDH2 inhibitor AG6780.

(**E**) Levels of enrichment (top panel) and IDH1 expression (bottom panel) in control fibroblasts or in the patient derived P893 fibroblasts transduced with control lentivirus or with the lentivirus harboring *IDH2*-specific shRNAs. Cells were collected at different time points after selection at 48 hours (lanes 2 and 6) and at 5 days (lanes 5 and 6). Note that none of the differences in this panel were statistically significant, in spite of the clear trend. This is likely due to the difficulty of measuring low D-2HG levels in the small percentage of transfected cells in these assays.

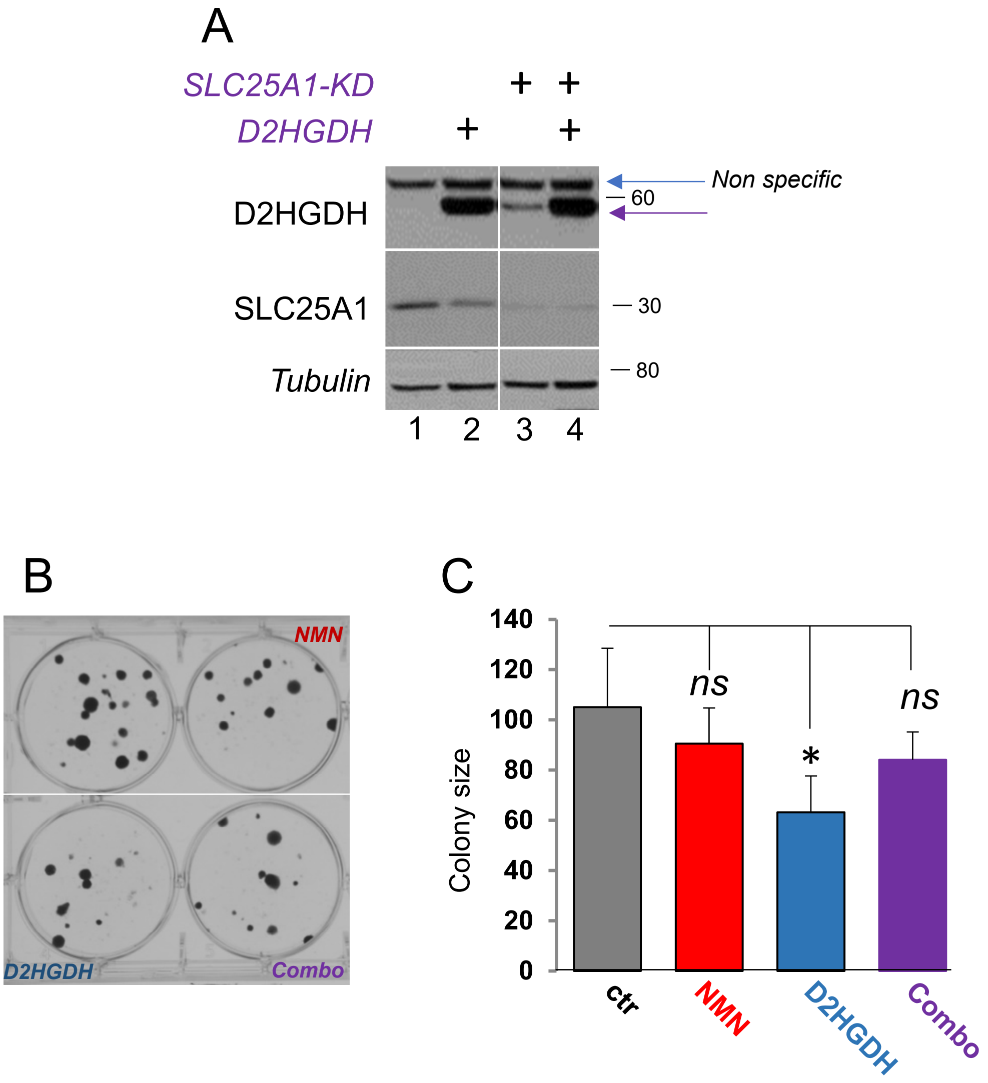

A
SLC25A1-KD
D2HGDH
+ +
+ +
D2HGDH
SLC25A1
Tubulin
Non specific
60
30
80
1 2 3 4
B
NMN
D2HGDH
Combo
C
Colony size
140
120
100
80
60
40
20
0
ns
*
ns
ctr
NMN
D2HGDH
Combo

**EV4. Effects of D2HGDH over-expression.**

(**A**) Immunoblot of H1299 cells over-expressing D2HGDH (lanes 2 and 4) in the presence or absence of the SLC25A1-KD (lanes 3,4). The levels of D2HGDH, SLC25A1 and Hsp70 are indicated. Arrows point to a non-specific band detected by the D2HGDH antibody.

(**B,C**) Representative images (**B**) and quantification (**C**) of colony forming ability in cells transduced with control lentirvirus.

**Supplemental Video 1.** Heart dysfunction in zebrafish embryos treated with 40 μM D-2HG at 36 hpf. The top embryo is treated, the bottom is control.
